# Direct and community-driven selection jointly drive body size evolution in harvested predator-prey systems

**DOI:** 10.64898/2026.03.01.708819

**Authors:** Théo Villain, Jean-Christophe Poggiale, Brandon Duquenoy, Nicolas Loeuille

## Abstract

Fisheries-induced evolution (FIE) affects multiple life-history traits, most notably body size. This evolutionary response is often examined at the single-species level and attributed to direct size-selective harvesting. However, fishing also targets other community members, reshaping trophic interactions and thereby modifying evolutionary constraints due to community changes. Here, we disentangle two forms of fishing-induced selection on body size - direct, arising from size-selective harvesting within species individuals, and indirect, emerging from fishing-induced changes in community structure - and investigate how their interplay shapes evolutionary trajectories. Using an adaptive dynamics framework within a predator-prey model, we show that (*i*) community destructuring can either amplify or dampen the effects of intraspecific size-selectivity on body-size evolution, (*ii*) predator evolution is primarily driven by direct selection, whereas prey evolution is mostly constrained indirectly by community structure. We then extend our analysis using a stochastic framework and show that (*iii*) prey evolution allows the evolutionary rescue of predators, underscoring the importance of community context. Our results demonstrate that eco-evolutionary feedbacks can profoundly alter both community structure and fishery yields, strengthening calls to incorporate evolution into ecosystem-based fisheries management.

## Introduction

Over the past few decades, fishing has been shown to induce changes in the life-history traits of exploited populations. Some of these changes are attributed to phenotypic plasticity and are reversible (Hendry, 2016), while others result from genetic changes (Kuparinen and Merilä, 2007). Fishing induces evolution of growth rates (Conover et Munch 2002; Enberg et al, 2011), size and age at maturity (Ernande et al, 2004), ultimately driving evolutionary body downsizing of exploited species (Olsen et al, 2004). These effects have been empirically documented (Barot et al., 2004; Carlson et al., 2007; Edeline et al., 2009, 2007; Grift et al., 2003; Olsen et al., 2004; Swain et al., 2007), experimentally demonstrated (Conover and Munch, 2002; Uusi-Heikkilä et al., 2015; Van Wijk et al., 2013), and supported by modeling approaches (Ernande et al., 2004; Jusufovski and Kuparinen, 2020). This growing body of research has led to broad recognition of the genetic basis of fisheries-induced evolution (FIE) (for reviews, see Ahti et al., 2020; Heino et al., 2015; Hočevar and Kuparinen, 2021; Palkovacs et al., 2012). Although there is ongoing debate regarding the rate of such evolutionary changes (Andersen and Brander, 2009; Audzijonyte et al., 2013a; Hilborn and Minte-Vera, 2008), it can sometimes occur rapidly (Olsen et al., 2004), on timescales comparable to those of ecological changes (Conover and Munch, 2002). Numerous studies now emphasize the importance of accounting for these effects in fisheries management (Conover and Munch, 2002; Garcia et al., 2012; Zhou et al., 2010), as they pose risks both to long-term yields (Conover and Munch, 2002) and to the ecological structure of communities (Palkovacs et al., 2012).

Fishing targets individuals based on their availability (Andersen et al., 2012; Salas et al., 2004) and specific traits of interest (Allendorf et al., 2008; Darimont et al., 2009). By selectively harvesting certain size classes (Allendorf et al., 2008), fishing increases the survival probability of individuals with contrasting phenotypes. Since body size is a heritable trait (Mousseau and Roff, 1987) and fishing-induced mortality is often intense (Olsen and Moland, 2011), evolutionary responses can occur (Audzijonyte et al., 2013a; Daufresne et al., 2009) and the direction of such changes depends on the differential selection pressure on body sizes (Lush 1937). These evolutionary effects have important consequences at the population level, as body size reflects a wide range of individual physiological processes (Ahti et al., 2020; Brown et al., 2004). It serves as a reliable proxy for reproductive success (Barneche et al., 2018) and intrinsic mortality (Kuparinen et al., 2023; White et al., 2013), with larger individuals typically experiencing lower mortality and higher reproductive output than smaller ones (Hixon et al., 2014). At the population scale, fisheries-induced evolution toward smaller body sizes threatens stock replenishment (Hutchings and Reynolds, 2004), reduces the economic value of individuals as prices are often size-dependent (Zimmermann and Heino, 2013), and ultimately undermines the long-term sustainability of fisheries (Conover and Munch, 2002; Hixon et al., 2014). Of major importance, body size also governs trophic interactions (Brose et al., 2006; Jennings et al., 2001), which in turn are central to the structuring of ecosystems (Bailey et al., 2010). At the community level, evolutionary reductions in prey body size have been shown to increase predation pressure and amplify the effects of fishing (Audzijonyte et al., 2013b), while evolutionary changes in predator size can trigger trophic cascades (Shackell et al., 2010) and destabilize entire ecosystems through increased population fluctuations (Kuparinen et al., 2016).

Fishing deeply alters marine communities (e.g., Daskalov et al, 2007, Frank et al, 2005) and drives changes in community species composition and the associated size structure (Bell et al, 2018). If FIE can generate significant ecological consequences at the community level, the reverse may also hold true. Community interactions shape fitness landscapes and fisheries-induced changes in community structure therefore influence evolutionary processes through eco-evolutionary feedback loops (Edeline and Loeuille, 2021; Loeuille, 2019; Palkovacs et al., 2012). For instance, predation is a key factor explaining prey life-history traits evolution (Day et al., 2002, Abrams 2000) and predator decline due to fishing can thus trigger cascading evolutionary effects on their prey (Palkovacs et al., 2011). Through density-dependent effects, predators can either drive a reduction in prey size (Heins et al., 2016) or lead to an increase in size (Reznick et al., 2001), in magnitudes similar to those caused by fishing (Reznick et al., 1990). Conversely, changes in prey densities can influence the evolution of predator body size (Edeline et al., 2007).

Eco-evolutionary feedback loops have been documented (Nilsson et al., 2019; Palkovacs et al., 2009; Palkovacs and Post, 2008), with selection induced by size-selective harvesting in a given stock sometimes constrained by community structure (Carlson et al., 2007; Edeline et al., 2007). However, none of the studies explicitly examine how fishing itself alters community structure which in turn generates additional selections on body size. Yet, the relative contributions of community-mediated (indirect) versus within-stock size-selective (direct) selection in driving fisheries-induced evolution remains largely unexplored (Edeline and Loeuille, 2021), despite the potential strong implications for fisheries management. Body size evolution in response to fishing may result from a synergy or antagonism between direct and indirect selection. When selections act in the same direction, their synergy can amplify the evolutionary response (Edeline and Loeuille, 2021), and rapidly favor certain phenotypes (e.g., slow-growing pikes during perch collapse in Edeline et al., 2007). Conversely, both selections can have antagonistic effects (Carlson et al., 2007; Edeline et al., 2007), potentially explaining cases where no visible phenotypic evolution occurs (*e*.*g*., Hilborn and Minte-Vera, 2008).

Evolution can theoretically rescue populations facing a disturbance by restoring their growth rate (evolutionary rescue; Gomulkiewicz & Holt, 1995; Carlson et al., 2014). However, it remains unclear whether this concept can be extended to communities and promote species coexistence (Loeuille, 2019). For instance, when predators are affected by a perturbation, prey evolution can prevent their extinction in a process known as indirect evolutionary rescue (Yamamichi & Miner, 2015). This has only been demonstrated when predators are the only impacted species. In the case of fishing, multiple species are simultaneously exploited, making such predictions challenging. FIE has been shown to limit stock replenishment compared to non-evolving populations (Conover & Munch, 2002). Within a community, if several species evolve in response to fishing, this could either improve their status or accelerate their decline (Loeuille, 2019). Understanding the contributions of different species to evolutionary outcomes at the community scale is therefore critical for assessing whether evolution can effectively maintain coexistence under exploitation and has important implications for fisheries management.

In this study, we aim to disentangle FIE into effects arising from fishing-induced changes in community structure (indirect selection, IS) and effects arising from size-selective harvesting within exploited species (direct selection, DS). To this end, we use a predator-prey model in which both trophic levels can be harvested. We explore different community destructurings by distributing fishing effort between prey and predators (changing interspecific selectivity, Figure 1C). Within each targeted species, specific size classes may be preferentially harvested (intraspecific selectivity, Figure 1B) and we evaluate different size-selective functions. In the first part of the study, we assume that evolution follows an adaptive dynamics process to isolate IS and DS. We first ask (*i*) to what extent IS affects FIE through the sole modification of community structure, and (*ii*) how IS and DS interact to shape body-size evolution and what ecological and economic consequences arise from this interplay. In the second part of the study, we move beyond the deterministic framework and investigate using a stochastic approach whether (*iii*) the concept of evolutionary rescue can be extended to the community scale.

**Figure 1:**
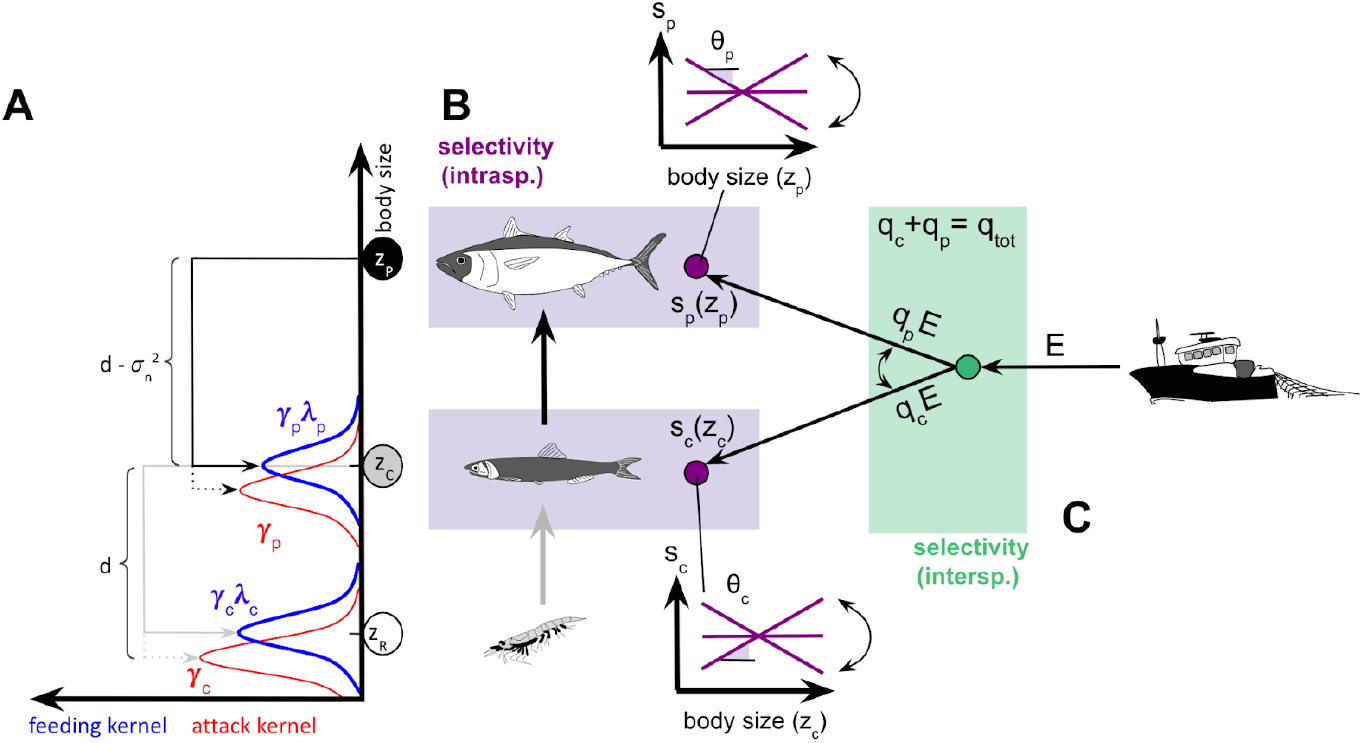
Trophic control of body size and fishing strategies. A) Representation of the community and associated trophic interactions. The preferred prey size of a predator is *d* smaller than its own size, but the predator maximizes resource acquisition when the prey is larger than this preferred size (difference of 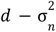), due to energetic transformation costs. B-C: The community is harvested with an effort *E* distributed between prey and predators (*q*_*c*_ and *q*_*p*_), representing interspecific selectivity (green, C). Within each stock, individuals are differentially harvested according to their size and the intraspecific selectivity (function *s*_*i*_ with slope θ_*i*_ purple, B).

We expect that the sole modification of community structure will lead to the evolution of both prey and predator species (*i*), but that the evolutionary changes will be more pronounced in prey species, as the impact of fishing on predator densities is greater than on prey densities. We also anticipate that both direct and indirect selection will either amplify or attenuate (*ii*) depending on fishing strategies, but that the magnitude of these effects will be greater in prey species due to (*i*), with significant consequences for yields. Finally, we hypothesize that evolution will favor the persistence of communities in the face of fishing, even when the associated ecological structures are severely degraded (*iii*).

## Methods

### 1) Ecological model and fishing

We consider a fished community composed of prey and predator compartments interacting within a simplified Lotka-Volterra model (eq.1 & 2):

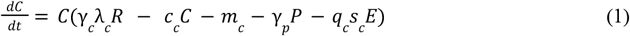

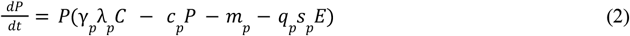

Where *C* and *P* represent prey and predator densities, respectively. Both species are subject to intraspecific competition (*c*_*c*_ and *c*_*p*_) as well as intrinsic mortality (*m*_*c*_ and *m*_*p*_). Preys consume an invariant resource *R* at an attack rate *γ*_*c*_, and convert the ingested energy into growth with an efficiency *λ*_*c*_. Analogously, predators attack prey at a rate *γ*_*p*_ and convert a fraction *λ*_*p*_ of the captured prey into their own growth (see Figure 1A). All these parameters depend on the body size *z* of individuals within species (Brose et al., 2006), defined as the natural logarithm of adult individual biomass (*z* = *log*(*biomass*)). Our model builds on previous approaches of body evolution, more specifically on Brännström et al (2011). Attack rates vary with the body size difference between species and reach a maximum when this difference equals *d* (gaussian variation, see Supplementary S1) (Brose et al 2006, Naisbit et al 2011, Li et al 2022), whereas conversion efficiency increases with the ratio of prey to predator (or ressource) body size (see details in S1). Finally, intrinsic mortality in both prey and predator compartments decreases with their size (Brown et al., 2004).

#### Interspecific selectivity of fishing

To account for the broad diversity of ecological consequences of fishing while allowing for comparisons across them, a total fishing effort *E* is distributed across trophic levels through *q*_*c*_ and *q*_*p*_, the respective catchabilities of prey and predators (*q*_*c*_ + *q*_*p*_ = *q*_*tot*_). Because *q*_*tot*_ is here fixed, when *q*_*c*_ is high, the fishery predominantly targets prey and only a few predators are captured, whereas *q*_*c*_ = *q*_*p*_ represents a balanced scenario. The allocation of effort defines the interspecific selectivity of the fishery (Figure 1C and BOX 1), and the ecological consequences of varying allocations constitute a first central dimension of the analysis.

#### Intraspecific selectivity of fishing

Within each of the two species, certain body-size classes are preferentially captured through the selectivity functions *s*_*c*_ and *s*_*p*_ and these functions modulate individual catchabilities. This pattern reflects not only the different techniques or size-based quotas that fishers may use to target individuals, but also potential spatiotemporal variability in body-size distributions. Because many scenarios could be construed, we explore the full range of possibilities by assuming that intraspecific size-selectivity functions can increase, decrease or show no variation with size (eq. 3):

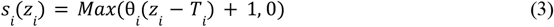

where *i* refers to the prey or predator compartment. If fishers target species *i* and focus on the size *T*_*i*_, individuals of size *z*_*i*_ = *T*_*i*_ are captured with a baseline efficiency of *s*_*i*_ (*z*_*i*_) = 1. In the stock, individuals of other sizes may then be captured more, equally, or less efficiently than the targeted size *T*_*i*_, depending on the slope θ_*i*_ of the intraspecific size-selectivity function. A positive slope θ_*i*_ implies that larger (smaller) individuals are more (less) efficiently captured, whereas a negative slope indicates the opposite pattern. For simplicity, we define the intraspecific size-selectivity of harvesting species *i* using the slope θ_*i*_ of these functions (Figure 1B and BOX 1). These slopes define a second main axis of investigation and their effects on eco-evolutionary dynamics are analysed later in detail.

A fishing policy is thus defined by the overall effort affecting the community (*E*), the interspecific selectivity (*q*_*c*_, *q*_*p*_), and the intraspecific selectivities (θ_*c*_, θ_*p*_) (see Figure 1). We assume that fishers do not adjust their practices even when stocks decline or species evolve in response to fishing, potentially reducing their catchability. While these assumptions are clearly simplifications, they may reflect the inherent inertia of fisheries, encompassing technical constraints, the difficulties of changing established practices or securing financing, regulatory limitations, and the fact that fishing strategies may also be constrained by the exploitation of other species (Schuch et al., 2021; Tudela & Short, 2005).

### 2) Approach and eco-evolutionary models

To investigate the effects of fishing on body-size evolution and its community-level consequences, we conduct a three-part study. In the first two parts, we use adaptive dynamics to assess the contribution of changes in community structure alone to body size evolution (IS), and then combine these effects with those of size-selective harvesting (IS + DS). In the third part, we explore whether evolution can promote species coexistence under fishing using a stochastic approach.

#### Adaptive Dynamics

We first use the adaptive dynamics framework to study the effects of fishing on the evolution of body size (Dieckmann and Law, 1996; Geritz et al., 1998; Metz, 1992). In short, mutants arise rarely (i.e., low mutation rate μ_*i*_ eq. 4) have a body size close to the resident population trait (i.e., small mutation amplitude *ν*_*i*_ eq. 4), and are initially rare compared to residents. Ecological equilibrium is therefore reached before the next mutation occurs, which allows ecological and evolutionary processes to be separated. Depending on the community, there can be substantial differences in the evolutionary potential of species (e.g., asymmetries in genetic variability). To account for these factors, we systematically investigate three evolutionary scenarios in the (first) two parts of the study embedded within the adaptive dynamics framework: (*a*) only the prey can evolve, (*b*) only the predators can evolve, and (*c*) both species coevolve. In each case, we analyze the consequences of fishing on size evolution under the assumption that prey and predator have reached a coevolutionary equilibrium prior to the onset of fishing. For simplicity, we describe below only the assumptions of model (*a*), while the descriptions of models (*b*) and (*c*) are provided in Supplementary S4.

Prey size (*z*_*c*_) evolution is assessed using the canonical equation of adaptive dynamics (eq. 4, Dieckmann and Law, 1996).

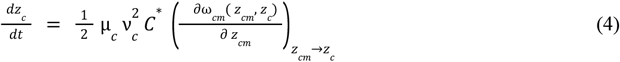

Where μ_*c*_ denotes the *per capita* mutation rate of prey, *ν*_*c*_ the mutation amplitude (see Table 1) and *C*^*^ the prey density at the ecological coexistence equilibrium with predators (see Supplementary S3&S4). ω_*cm*_ is called the invasion fitness (Metz et al 1992), defined by the rate of variation of the density of rare mutants of body size *z*_*cm*_ in a resident population of trait *z*_*cm*_ (eq. 5):

**Table 1:** Variables and parameters used in the eco-evolutionary models. Values are given for the adaptive dynamics model (AD), the stochastic model with initial regular (SRP) or inverted pyramid (SIP). The intensity of species interactions are then calibrated using size-based relationships (see Supplement S1). Initial size values at eco-evolutionary equilibrium in single evolution experiments (1) or in stochastic co-evolution (2). (3) These parameters are investigated to balance fishing selections. nd: not defined.

| Variables | Biological meaning | Unit | Value (AD / SRP / SIP) |
| --- | --- | --- | --- |
| $C$ | Prey density | ind m<sup>-2</sup> | - / - / - |
| $P$ | Predator density | ind m<sup>-2</sup> | - / - / - |
| $z_c$ | Prey size | log(kg) | 5.75<sup>(1)</sup> / 6.16<sup>(2)</sup> / 6.61<sup>(2)</sup> |
| $z_p$ | Predator size | log(kg) | 10.40<sup>(1)</sup> / 10.81<sup>(2)</sup> / 11.26<sup>(2)</sup> |
| $t_{ext}$ | Time before predator extinction | t | nd / - / - |
| Parameters | Biological meaning | Unit | Value (AD / SRP / SIP) |
| $\gamma_0$ | Per capita basal attack rate | ind<sup>-1</sup>m<sup>2</sup>t<sup>-1</sup> | 10 / 10 / 40 |
| $\lambda_0$ | Basal conversion efficiency | - | 0.3 / 0.3 / 0.45 |
| $m_0$ | Basal mortality rate | t<sup>-1</sup> | 0.1 / 0.1 / 0.1 |
| $d$ | Predator-prey optimal feeding distance | log(kg) | 5.65 / 5.65 / 5.65 |
| $z_r$ | Resource size | log(kg) | 1 / 1 / 1 |
| $e_0$ | Conversion parameter (g to kg) | - | 10<sup>-3</sup> |
| $c_c$ | Per capita intraspecific competition rate of preys | ind<sup>-1</sup>m<sup>2</sup>t<sup>-1</sup> | 1 / 1 / 1 |
| $c_p$ | Per capita intraspecific competition rate of predators | ind<sup>-1</sup>m<sup>2</sup>t<sup>-1</sup> | 1 / 0.1 / 0.1 |
| $E$ | Fishing effort | effort | 1 / 250 / 5800 |
| $q_{tot}$ | Total catchability | t<sup>-1</sup>effort<sup>-1</sup> | 1 / 1 / 1 |
| $q_c$ | Prey fishing catchability (independent of size-selective effects) | t<sup>-1</sup>effort<sup>-1</sup> | [0, $q_{tot}$]<sup>(3)</sup> / 0.5 / 0.5 |
| $q_p$ | Predator fishing catchability (independent of size-selective effects) | t<sup>-1</sup>effort<sup>-1</sup> | [0, $q_{tot}$]<sup>(3)</sup> / 0.5 / 0.5 |
| $\theta_c$ | Slope of prey fishing size-selectivity | log(kg)<sup>-1</sup> | [- 1, 1]<sup>(3)</sup> / 0 / 0 |
| $\theta_p$ | Slope of predator fishing size-selectivity | log(kg)<sup>-1</sup> | [- 1, 1]<sup>(3)</sup> / 0 / 0 |
| $R$ | Basal resource density | ind m<sup>-2</sup> | 5 E3 / 5 E5 / 5 E5 |
| $\mu_c$ | Prey mutation rate | t<sup>-1</sup> | 10<sup>-4</sup> / [0, 10<sup>-1</sup>] / [0, 10<sup>-1</sup>] |
| $\mu_p$ | Predator mutation rate | t<sup>-1</sup> | 10<sup>-4</sup> / [0, 10<sup>-1</sup>] / [0, 10<sup>-1</sup>] |
| $v_c$ | Prey amplitude of mutation (standard deviation) | log(kg) | 0.15 / 0.15 / 0.15 |
| $v_p$ | Predator amplitude of mutation (standard deviation) | log(kg) | 0.15 / 0.15 / 0.15 |
| $\tau$ | Time step in stochastic simulations | t | nd / 1E - 4 / 1E - 5 |

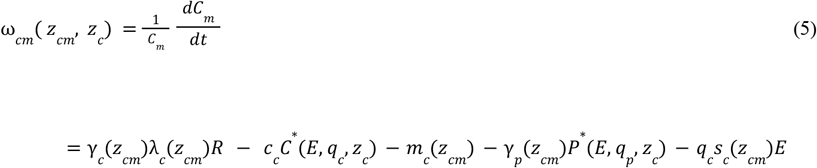

Where *P*^*^ denotes the predator density at the ecological coexistence equilibrium with prey (see Supplementary S3&S4). Predator body size is assumed not to evolve in this case (*a*). According to eq. 4, the evolution of prey size depends on the ability of nearby mutants to invade the system defined by residents. Evolution proceeds if the population persists (*C*\* ≠0) and as long as proximate sizes confer a competitive advantage (∂ω_*cm*_ ≠0). When the selection gradient (eq. 6) is positive (negative), it means that a mutant of larger size has better (worse) fitness compared to the residents, and this is reflected in the canonical equation (eq. 4) by the increase (decrease) in size through invasion. Harvesting the community through interspecific (*q*_*p*_, *q*_*c*_) and intraspecific (*s*_*c*_) selectivities may affect prey evolution both indirectly, via changes in community structure (changes in *P*^*^ in third term of eq. 6), and directly, through selection on prey body size (fourth term of eq. 6):

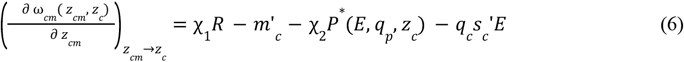

where

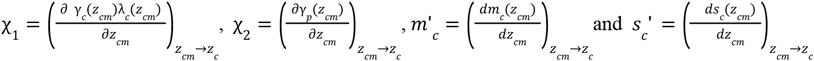

Depending on the sign of the third and fourth terms of eq. 6, DS and IS may either act in the same or in opposite directions, thereby influencing body size evolution. Once the selection gradient vanishes, the evolution of prey size stops and reaches the evolutionary singularity 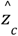.

#### Model analysis

In the first part, we examine whether fishing-induced modifications in community structure can drive body size evolution through IS. To this end, we isolate the effects of density variations by removing intraspecific selectivity (*s*′_*i*_ = θ_*i*_ = 0 in eq. 3). In the second part, we investigate how DS and IS interact in response to fishing, and the consequences of these interactions for both community structure and fishery yields. In particular, we aim to identify the conditions under which these forces combine synergistically to amplify evolutionary responses, or alternatively counteract each other with antagonistic effects. Community structure at eco-evolutionary equilibrium is then described by prey and predator densities (*Ĉ* and 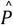), their respective biomasses (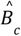 and 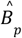, eq. 7), the total biomass (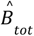, eq. 8), and the Shannon index (*Ĥ*, eq. 9). Although this index is most commonly applied in species-rich communities, here it serves as a proxy for maximum trophic height or for the shape of the abundance pyramid. Specifically, values of *Ĥ* close to 0 indicate that predators are present but at very low density (corresponding to low trophic height and a bottom-heavy pyramid), whereas maximal values occur when predators and prey are equally abundant (reflecting greater trophic height and a more homogeneous pyramid).

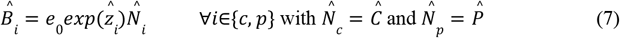

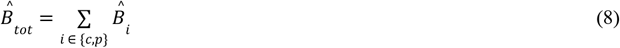

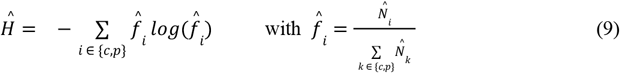

The yields *Ŷ* are calculated at the eco-evolutionary equilibrium as the total biomass harvested by fishing (eq. 10).

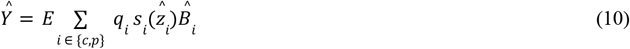

#### Stochastic simulations of the eco-evolutionary dynamics

In the final part, we examine whether body-size evolution can promote community coexistence under intense fishing pressures, thereby extending the concept of evolutionary rescue to a community-level perspective. To study population persistence over time, we assume that evolution can occur outside of ecological equilibrium, *i*.*e*., ecological and evolutionary processes operate on the same timescale. Accordingly, we perform stochastic simulations using the tau-leap method (Gillespie, 2001) in a coevolving system under different combinations of prey and predator mutation rates. Time is discretized into intervals of length τ. Within each interval, population births and deaths are sampled from Poisson distributions based on the states at the beginning of the interval. Mutants arise from the births, with traits drawn from a Normal distribution centered on the parental trait and exhibiting an average 5% deviation. To evaluate the effect of evolution on community persistence, we first simulate 10^4^ time steps without fishing, starting from monomorphic prey and predator populations whose sizes correspond to the deterministic eco-evolutionary equilibrium. The system is then subjected to exploitation at an intensity that drives predator extinction in the absence of evolution. We compare predator extinction times (*t*_*ext*_ in Table 1) with and without coevolution using a logarithmic ratio 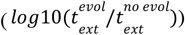. This analysis is conducted for communities with either regular (SRP) or inverted (SIP) biomass pyramids.

##### BOX 1 : Definitions

*Intraspecific selectivity:* means implemented to preferentially target certain size classes among individuals belonging to the same species. This is represented by an affine function of body size *s*_*i*_(*z*_*i*_) with slope θ_*i*_, which is often referred to as intraspecific selectivity for convenience in the text.

*Interspecific selectivity:* means implemented to preferentially harvest certain species within a community. This is represented by the distribution of the total fishing effort *E* among species and scales with (*q*_*c*_,*q*_*p*_).

*Direct selection:* in a fitness gradient related to the size of a species, it is an evolutionary force that results from fishing size-selectivity. The intensity of this selection depends on intraspecific selectivity, interspecific selectivity, and the effort used to fish the community.

*Indirect selection*: in a fitness gradient related to the size of a species, it is an evolutionary force that results from the modification of ecological constraints on size, linked to changes in species abundances. The intensity of this selection depends on intraspecific selectivity, interspecific selectivity, and the effort used to fish the community.

## Results

### A) Indirect selection has a stronger impact on prey evolution due to constraints on fitness

To understand how IS can drive body-size evolution, we examine the evolutionary implications of community structure changes under fishing in the absence of DS (θ_*i*_ = 0, eq. 3). This analysis is carried out for the three evolutionary scenarios (models *a, b*, and *c*). We show that prey evolution is highly sensitive to IS (model *a*, Figure 2A), whereas predator evolution is almost unaffected (model *b*, Figure 2B). When both species coevolve in response to community perturbations caused by fishing, prey evolution drives predator evolution (model *c*, Figure 2C).

**Figure 2:**
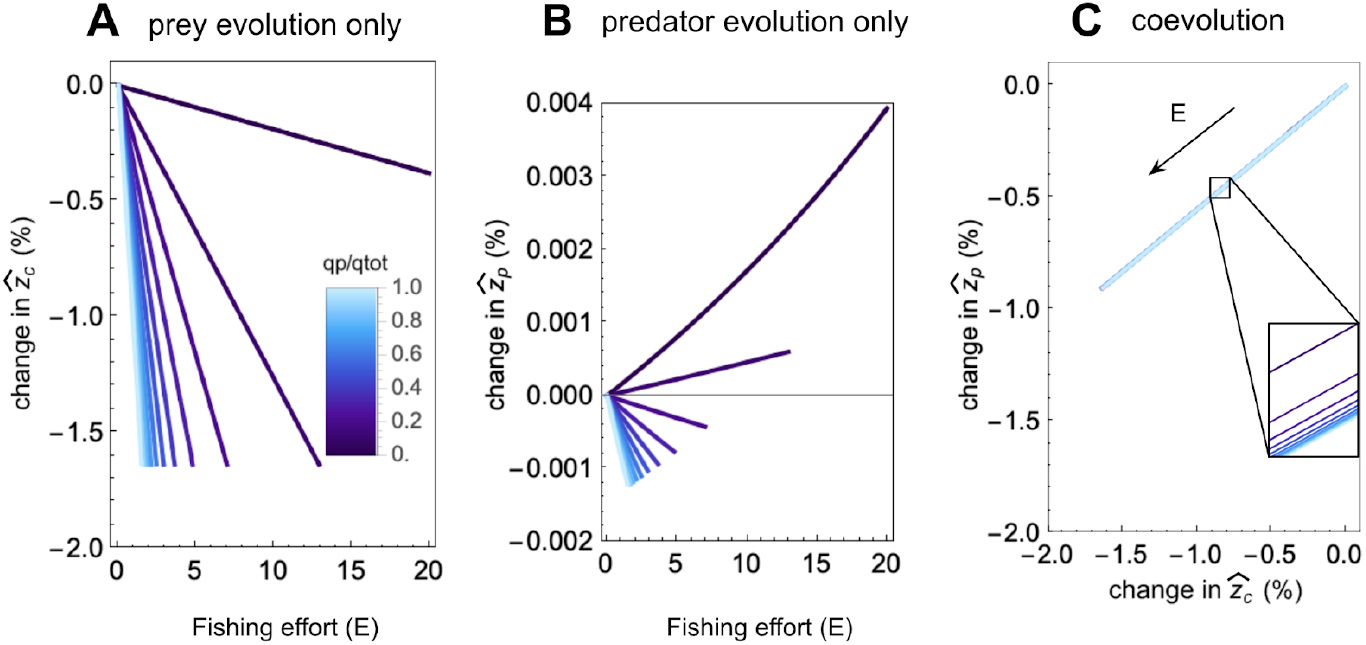
Evolution of prey and predator sizes in response to exploitation. A: Evolution of prey size in response to fishing effort under different strategies, with prey being the only species able to evolve. A-B-C: Fishing strategies are represented by shades of blue. Light blue: *q*_*p*_ = *q*_*tot*_,*q*_*c*_ = 0; dark blue: *q*_*c*_ = *q*_*tot*_,*q*_*p*_ = 0. B: Evolution of predator size when they are the only species able to evolve. C: Co-evolution of prey and predator sizes. Sizes are normalized to those obtained in the absence of fishing.

These results are linked to fitness constraints: in the absence of fishing, prey body size is determined by the trade-off between resource acquisition and predation-induced mortality (eq. 6). By decomposing the impact of fishing on predators, it becomes clear that predator abundance is always negatively affected, since they both compete with fishers for prey and are themselves subject to capture (eq. 11):

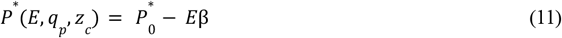

With 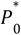 the predator density without exploitation and 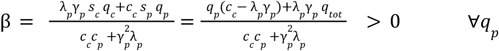, with *s*_*i*_ = 1 since θ_*i*_ = 0

Thus, eq. 6 can be rewritten by distinguishing the components independent of fishing pressure (first term, with exponent *E* = 0, eq. 12) from those that depend on fishing effort (second term):

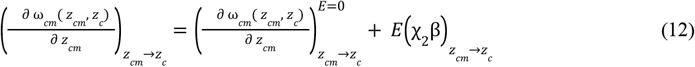

It is possible to show that the first term is a decreasing function (non-invasibility criterion, Metz et al., 1992) and that χ_2_ < 0 when prey and predator body sizes are initially at their long-term eco-evolutionary equilibrium (for other situations see Figure S7). Thus, eq. 12 shows that when predator density decline is substantial, it induces pronounced evolutionary decreases in prey body size (with a shift of about 1%, Figure 2A), driving prey toward sizes better adapted to resource acquisition (for a formal demonstration, see Supplementary S5 and eq. A18). Note also that when (*c*_*p*_ − *λ*_*p*_*γ*_*p*_) > 0 (i.e. competition dominates over predation, a bottom-up community), the magnitude of prey evolution is maximal when only predators are targeted (*q*_*p*_ = *q*_*tot*_).

Conversely, since they are not subject to predation, predator body size is primarily constrained by their capacity to acquire resources, i.e., by prey size. When preys do not evolve in size (model *b*), predators show only minimal evolutionary responses to fishing (on the order of 10^−3^%, Figure 2B). Although marginal, the impact of fishing on prey densities can either increase or decrease predator size (Figure 2B and Supplementary S6). This difference arises because prey densities may increase (*e*.*g*., when fishing strongly targets predators) or decrease (e.g., when fishing focuses on prey). By contrast, when both species coevolve (model *c*), prey-size evolution drives predator evolution, which then follows the size optimum for resource consumption (Figure 2C). Stronger predator depletion (high *q*_*p*_, shown in light blue in Figure 2C) leads to stronger evolutionary shifts in both species (see zoomed inset).

Thus, altering community structure changes the strength of trophic interactions and modifies constraints on body size. In response, prey evolution may be substantial and drives predator evolution when both species coevolve.

### B) Combinations of IS and DS have contrasting effects on body-size evolution

We now examine how IS and DS interact by incorporating size selectivity within each species. First, we aim to understand the consequences for evolution (Figure 3), and then how these effects feedback on community structure and fishery yields (Figure 4).

**Figure 3:**
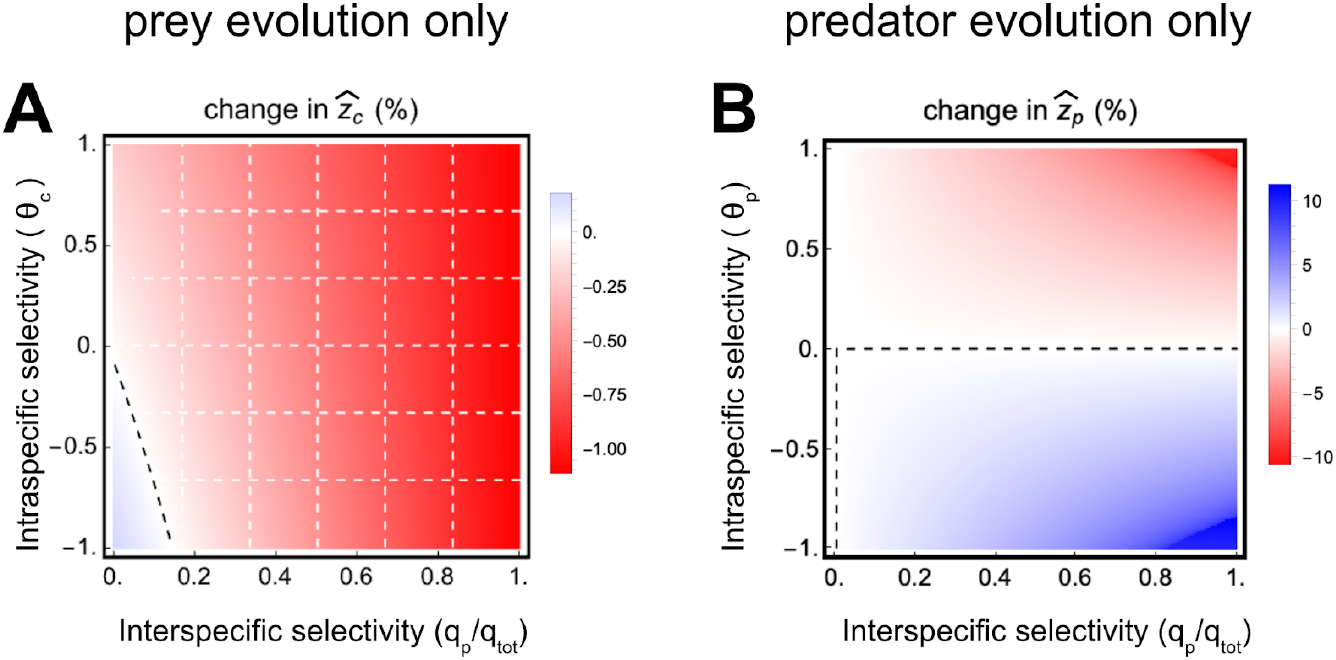
Evolution of sizes as a function of intraspecific and interspecific selectivity when only one species evolves. Depending on the nature of the fishing pressure, the size 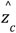 of preys (A) or 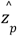 of predators (B) may either increase (in blue) or decrease (in red) due to evolution. The contribution of IS to size evolution can exceed that of DS (dashed white grid). DS and IS can offset each other, leading to the visible lack of size evolution (dashed black line). The fishing effort equals *E* = 1.

**Figure 4:**
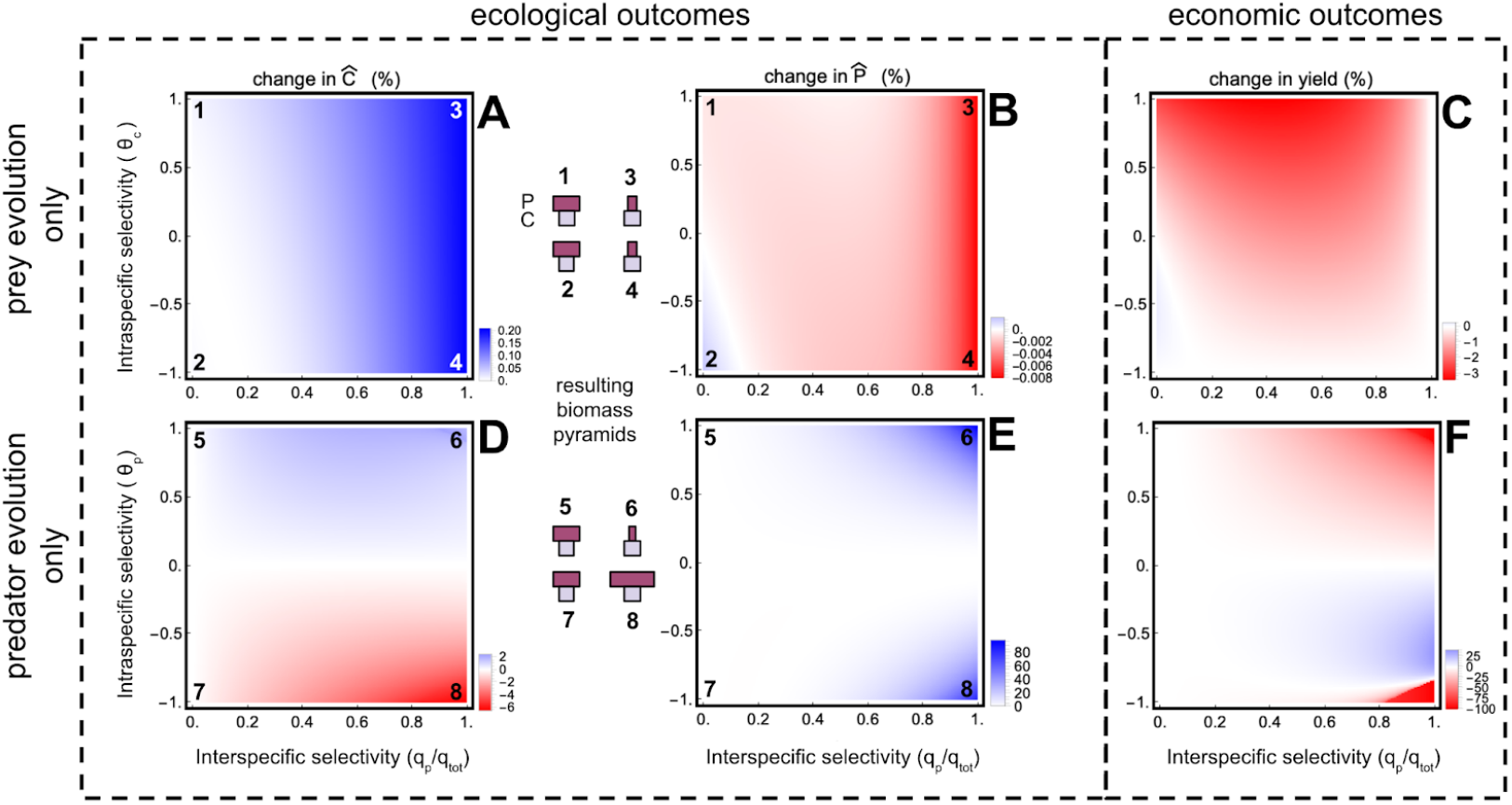
Consequences of size evolution on fishing yields and community structure when a single species evolves. Depending on the nature of the fishing pressure, prey densities (*Ĉ*), predator densities 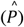, and yields can either increase (blue) or decrease (red) compared to a situation without evolution. The fishing effort equals *E* = 1. A-B-C: only preys evolve. D-E-F: only predators evolve.

#### 1) The evolution of sizes depends on the trophic position of the evolving species

IS and DS can either act in the same direction, amplifying body-size evolution, or in opposite directions, limiting it (Figure 3). Based on the fitness gradient, we detail the reasoning only for prey evolution (model *a*, eq. 13), but the logic is strictly analogous for the other models. While in absence of DS, changes in community structure induce prey body-size evolution in only one direction (Figure 2A), accounting for DS allows fishing to cause increases, no change, or decreases in prey body size (eq. 13 and Figure 3A). When large prey are targeted (*i*.*e*., θ_*c*_ > 0), DS and IS reinforce each other (since χ_2_ < 0), and fishing drives strong evolution toward smaller body sizes (Figure 3A, in red).

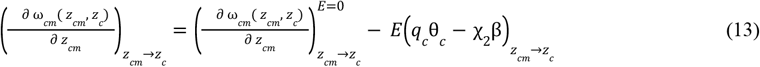

Conversely, when small prey are mostly targeted (*i*.*e*., θ_*c*_ < 0), IS and DS act in opposition, dampening body size evolution. Note that if prey and predator body sizes were not at the eco-evolutionary equilibrium prior to fishing, alternative outcomes could occur (see Figure S10), but the underlying mechanism remains unchanged. While in most fishing conditions IS has a stronger effect than DS (indicated by white hatching in the figure), this pattern is reversed when fishing effort is primarily directed at prey (small *q*_*p*_, left of the panel 3A). Under these conditions, the effect of DS is stronger (since |*q*_*c*_ θ_*c*_| is higher) whereas the effect of IS is weaker (since β is lower). IS can then counteract the effects of DS, leading to an increase in prey body size due to fishing (Figure 3A, in blue).

Similarly for predators (Figure 3B), the two selection pressures can act in the same direction or oppose each other, but here DS primarily governs body size evolution (no white dashed areas). Previous results show that changes in community structure have little effect on predator evolution, because it mainly depends on prey size (Figure 2). Consequently, when predators are the only evolving species (model *b*), IS is weak and their evolution is largely driven by DS (Figure 3B, with colors varying along the y-axis). Concentrating fishing on predators (right of the panel) slightly increases the amplitude of evolution, as the selection pressure is stronger (*q*_*p*_ *s*_*p*_ *E*, eq. 2). Note also that the evolutionary amplitudes are larger for predators (up to 10%) than for prey (up to 1%). Although these effects are difficult to fully explain, they may result from a strong trade-off affecting prey size between resource acquisition and mortality due to predation (i.e., a steep fitness gradient), whereas this trade-off is weaker for predators. In such a scenario, the same intensity θ_*i*_ of selection would be more costly for prey than for predators, which could account for the observed differences.

Interestingly, for both prey and predators, when the two selection pressures have equal magnitude but act in opposite directions (*e*.*g*., *q*_*c*_ θ_*c*_ = χ_2_ β for prey evolution but see Supplementary S6 for complete developpement), no body size evolution would be observed (eq. 13 and Figure 3, black dashed lines). This effect is independent of the total fishing effort applied to the community (eq. 13) and depends solely on its distribution between the two compartments. Paradoxically, this phenomenon cannot occur when a species is not subject to fishing (*e*.*g*., *q*_*c*_ ≠0).

#### 2) The evolution of predators has stronger consequences on yields and the community

We now examine the consequences of the combined IS and DS on community structure (*Ĉ* and *Ĉ*) and yields (*Ŷ*, eq.10), by comparing the outcomes of identical fishing scenarios with and without evolution. Ecological and economic effects of evolution are greater when fishing targets predators, and even more pronounced when predators themselves evolve (Figure 4). Ecological outcomes primarily depend on interspecific selectivity when prey evolve, whereas they mainly depend on intraspecific selectivity when predators evolve (Figure 4ABDE). These effects arise because body size evolution in each species directly translates into changes in their densities. When preys evolve (model *a*, Figure 4AB), their densities increase slightly (by ∼ 10^−1^%), and this effect is stronger when fishing targets predators, i.e. when IS is strong. Surprisingly, predator densities are only marginally affected by prey size evolution (Figure 4B, changes of about 10^−3^%). Conversely, when predators evolve, prey densities are more strongly affected (Figure 4D, ∼ 2%), with the direction of change depending on intraspecific selectivity (θ_*p*_) since predator evolution is primarily driven by DS. When fishing drives predators to evolve towards larger body sizes, prey densities are further reduced, as they are more heavily consumed (Figure 1A, and analogously with χ_2_ < 0). Predator evolution also results in a marked increase in predator densities (Figure 4E), mainly because, by evolving in response to DS, they eventually become less susceptible to capture. The combination of these effects leads to a system in which predator biomass largely dominates (Figure 4, point 8) and homogenizes community structure (Shannon index, Figure S12), compared to the same fishing scenario where only prey evolve (Figure 4, point 4).

Regarding fishing yields, we observe that they are often reduced by FIE (Figure 4CF, red dominates), mainly because body size evolution can make species less susceptible to capture. Interestingly, effects of prey evolution on yields are mostly linked to direct selection targeting large individuals (red area in upper panel C), while effects of predator evolution involve an interaction between direct and community selection (large effects in the top right and lower right of panel F). Prey evolution has a negative but limited effect on yields, as prey become more abundant but smaller, which translates into lower biomass (Figure S11) and reduced catchability. Predator evolution, by contrast, has a much stronger impact on yields (Figure 4E). When predators evolve towards smaller sizes, yields can decline by up to 40%, even though predator densities increase (Figure 4E), leading to a system dominated by prey biomass (Figure 4, point 6). Conversely, when intraspecific selectivity targets smaller sizes (θ_*p*_ < 0), yields can increase by nearly 40% (Figure 4F, blue), because predators become both larger and more abundant (Figure 3B and point 8). Once again, these large changes in yields result from the strong evolutionary responses of predators driven by DS (Figure 3B). Note that evolution may lead to sizes that escape fishing (individuals are no longer captured when 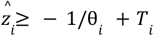). Since evolution in predators is significant and stronger when it favors larger sizes than smaller ones (Figure 3), this situation is more likely to occur when fishing targets small individuals. This scenario is not very realistic, as fishers never adjust their catches, but it poses a risk of losing 100% of the yield, since predators may become completely uncatchable (Figure 4). Thus, targeting smaller predators increases yields but may eventually lead to their decline if selection is too strong.

In coevolution scenarios (model *c*, Figure S13,S14,S15), the consequences for yields and ecological structure remain similar to the case where only predators evolve (Figure 4DEF) and are therefore much more pronounced than when only prey evolve (Figures S12,S13). Evolution in both species is broadly similar to that observed in models *a* and *b*, although prey evolution can be slightly higher. Thus, targeting predators has stronger impacts on evolution, ecological structure and yields (Figure S13,S14,S15). When *q*_*p*_ is high, fishing primarily targeting predators drives stronger prey size evolution, since prey size is mainly sensitive to IS and responds to changes in predator densities (Figure S13). This fishing strategy also has a greater impact on predator size, as its evolution is primarily driven by DS, which is stronger when *q*_*p*_ is high. Evolution toward larger predator sizes can increase fishing yields but may also lead to zero yields when sizes become too large to be caught (Figure S15). Additionally, it is possible that neither species evolves in size in response to fishing, with the conditions for this to occur being a combination of those required for each species in the case of individual evolution (*e*.*g*., *q*_*c*_ θ_*c*_ = χ_2_ β for prey evolution, see Supplementary S9).

Thus, when both forces are combined, it is clear that prey and predators respond to different selection pressures. Prey evolution is primarily driven by IS, whereas predator evolution is mainly driven by DS. Fishing concentrated on predators therefore has a greater impact on evolution and its consequences for community structure and fishing yields. Interestingly, whereas in the absence of DS, body size coevolution was mainly governed by prey evolution, the presence of both DS and IS shifts the driver of size coevolution to predator evolution.

### C) Prey evolvability is crucial for the evolutionary rescue of the community

Finally, we aim to assess whether the concept of evolutionary rescue (Gomulkiewicz and Holt, 1995) can be extended to the community level. Predicting coexistence through evolutionary rescue when both species are subject to fishing is not straightforward (Loeuille, 2019). This concept requires ecological and evolutionary dynamics to operate on similar timescales. To explore this extension, we move beyond adaptive dynamics - which assumes a separation of timescales - and examine the outcome of coevolution on predator extinction in a system where eco-evolutionary dynamics are modeled stochastically (Figure 5).

**Figure 5:**
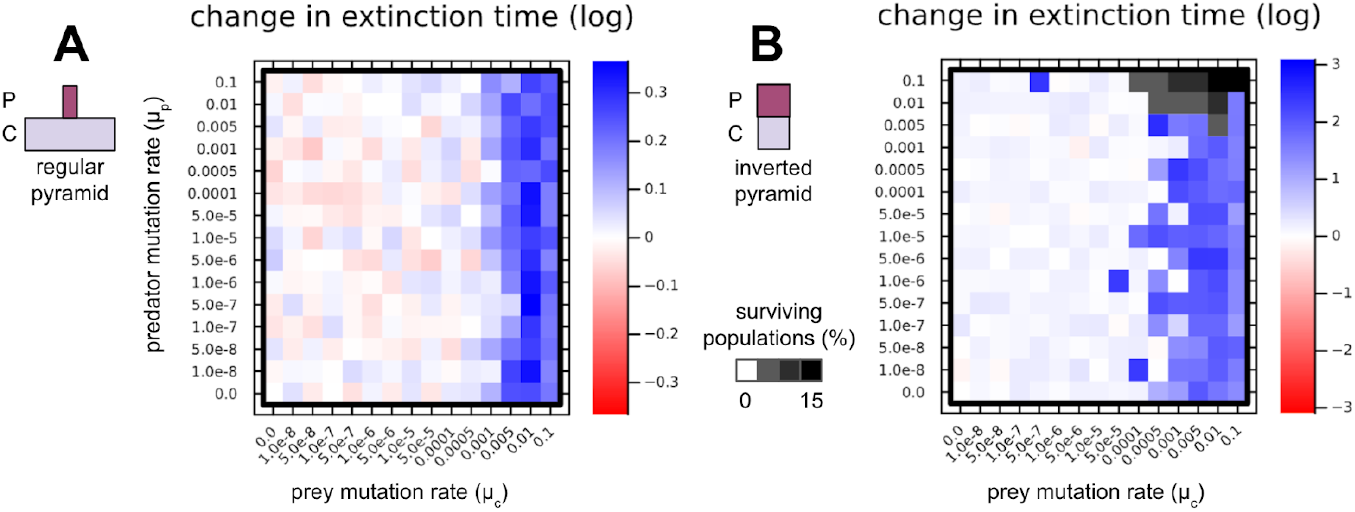
Effect of evolution on predator extinction time under intense fishing pressure. Fishing pressures are calibrated to ultimately lead to predator extinction in the absence of evolution.The color scale represents the logarithmic ratio of predator extinction times with 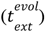 versus without 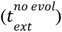 evolution 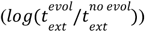. Blue (red): evolution allows predators to persist longer (shorter) compared to a scenario without evolution. We compare the effect of different mutation rates for prey size (μ_*c*_) and predator size (μ_*p*_) within a stochastic model, considering either a normal density pyramid (A, *n* = 100) or an inverted pyramid (B, *n* = 20). When some populations persist despite fishing, we record their proportion among all simulations and overlay the corresponding grayscale. A-B: fishing effort is equally distributed between prey and predators (*q*_*p*_ = *q*_*c*_ = 0. 5) with no intraspecific selectivity (θ_*c*_ = θ_*p*_ = 0). 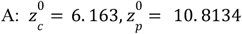, with a fishing pressure of *E* = 250. 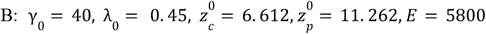 Other parameters used are the default values.

At the community level under fishing, evolutionary rescue is possible, but the contributions of different species are unequal, with prey evolution being critical for system persistence while predator evolution matters less (Figure 5). Evolutionary rescue is more likely when mutation rates are high (Carlson et al., 2014). We would therefore have expected that evolution in both prey and predators contributes to community maintenance (translating into blue regions above the diagonal in a figure like Figure 5). However, our results indicate that it is primarily prey evolution (starting from μ_*c*_ = 10^−3^, Figure 5A) that enables predator survival and thus constitutes an indirect evolutionary rescue. In this situation (balanced fishing and θ_*i*_ = 0), predator decline benefits prey through the release of predation pressure. Higher prey densities enhance their evolvability, and under strong mutation rates this allows predators to persist longer, likely because prey densities are higher at smaller sizes. Studying the case of a more homogeneous pyramid (stronger top-down control, Figure 5B) supports this hypothesis, and lower mutation rates in prey compared to Figure 5A allow predator persistence due to stronger effect of predation release. In communities with this ecological structure, predator evolution is not critical but does increase the magnitude of evolutionary rescue (shaded areas, Figure 5B), as predators persist until the end of the simulation.

We finally observe that coevolution in response to fishing also leads to higher cumulative yields (from the start of the perturbation to eventual predator extinction) compared to a scenario without evolution (Figure S14). However, because yield variations are strongly correlated with predator persistence, these higher yields are mainly due to the community being maintained for a longer period and thus available for harvest, rather than intrinsic effects of evolution on biomass.

## Discussion

In this study, we showed that the community context is crucial for understanding fisheries-induced evolution. Our results suggest that (*i*) modifying community structure can either amplify or mitigate the effects of intraspecific selectivity on body size evolution. Even in the absence of any regulation targeting the size of harvested individuals within a species (DS), harvesting predators alone alters trophic constraints and drives prey size evolution. These effects vary depending on how fishing effort is distributed within the community. When intraspecific selectivity is taken into account, IS may act in the same direction as DS and thus accentuate body size evolution. Conversely, the two selection types may act in opposite directions, attenuate each other, and may even result in no visible evolutionary change. The second key finding of this study is that (*ii*) at the community level, different trophic levels respond heterogeneously to fishing-induced selection. Prey are particularly sensitive to IS because of the trade-off between resource acquisition and vulnerability to predation, whereas predators are not subject to this trade-off, which makes them mainly vulnerable to DS. These evolutionary effects have direct consequences on community structure and yields, with impacts becoming stronger when predators are targeted and able to evolve, since predator harvesting amplifies both IS in prey and DS in predators, thereby explaining these results. We find that the community-level outcomes are contrasted, while yields are often reduced by these FIEs. Finally, (*iii*) our results highlight the importance of considering community context to explain persistence under high fishing pressure. Predator evolution alone is often insufficient to cope with strong disturbances, whereas prey evolution allows for persistence through a mechanism of indirect evolutionary rescue. Overall, our findings underline the need to account for evolution beyond the scale of a single managed stock, both for its evolutionary consequences at the community level and for its repercussions on ecological structure and fishery yields.

Size evolution in response to fishing has been demonstrated in numerous studies (Olsen et al. 2004, Ernande et al. 2004, Conover and Munch, 2002, Uusi-Heikkilä et al. 2015) but is often interpreted as the result of size-selective fishing (Swain et al. 2007). Our study, however, highlights the importance of community context in understanding size evolution. By altering density ratios, fishing affects the fitness constraints of individuals and thus directs size evolution. We describe these effects as indirect selection on body size by fishing rather than as natural selection. Some authors distinguish natural from artificial selection (Edeline and Loeuille, 2021, Edeline et al. 2007, Edeline et al. 2009, Carlson et al. 2007, Olsen and Moland, 2011) because they consider natural selection to be relatively independent of artificial selection. Moreover, this terminology often encompasses physiological (Carlson et al. 2007, Edeline et al. 2007) or behavioral components (cannibalism in Edeline et al. 2007 and risk-taking in Olsen and Moland 2011). In our case, fishing directly modifies the fitness landscape associated with interactions, generating size-selective pressures, which is why we prefer the terms direct and indirect selection. Nevertheless, our results align with the idea that size-selectivity can either counteract or reinforce other forces, thus dampening or amplifying size evolution due to fishing.

We showed that IS predominantly reduces prey body size, while having limited effects on predator evolution. For prey, these effects depend on the initial distribution of body sizes within the community, because vulnerability to predation follows a dome-shaped relationship (Fuiman & Magurran 1994; Edeline et al. 2007). At eco-evolutionary equilibrium, predators optimize resource acquisition and are consequently larger than the size maximizing their attack rate; their decline thus drives a reduction in prey body size. When predators are initially smaller, IS drives the opposite outcome and favors larger prey (Figure S7). Making a parallel between fishing and predator presence, these results echo observations of morphological traits with and without predators. For instance, Reznick et al. (2001) show that higher predator abundance favors the evolution of larger prey, and Palkovacs et al. (2011) experimentally demonstrated that predator removal directs prey evolution towards phenotypes specialized in resource acquisition. Regarding predator evolution, our results show that when predators are the only ones allowed to evolve, IS is very weak but can act in either direction. These effects may be consistent with Edeline et al. (2007), who showed that fewer prey promote the evolution of phenotypes with slower growth rates, in contrast to scenarios with abundant prey. In their study, however, it remains unclear whether this evolution leads to larger or smaller predators, making it difficult to directly relate our results to the direction of evolution in their maturation rates.When the whole community evolves together, our results suggest that the effects of IS on prey cascade onto predator evolution, leading to a reduction in predator body size. These observations can be related to the bioenergetic analysis of Shuter et al. (2015), in which smaller prey lead to smaller predators.

Finally, regarding the direction of size evolution, our findings align with previous studies. Selective fishing targeting larger phenotypes drives evolution toward smaller sizes, while the opposite occurs when fishing targets smaller phenotypes (Conover and Munch 2002, Uusi-Heikkilä et al. 2015, van Wijk et al. 2013). However, we demonstrate that the apparent absence of evolutionary change can result from the opposing effects of direct and indirect selection, regardless of the fishing effort applied. Our results thus contribute to the ongoing debate on the rate of fishing-induced evolution (Hilborn and Minte-Vera 2008, Andersen and Brander 2009, Audzijonyte et al. 2013) and suggest that these differences may be particularly influenced by community-level disturbances.

It is challenging to identify clear eco-evolutionary feedback loops (EEFLs) like those described by Palkovacs and Post (2008) in our study, as our results are obtained at eco-evolutionary equilibrium. Nevertheless, our study highlights not only the importance of the ecological effects of fishing on fisheries-induced evolution (FIE, as previously discussed) but also the reverse process. We show that FIE related to predator fishing has a greater impact on community structure than FIE associated with prey fishing. This is not surprising, given that predators play a key role in structuring community dynamics (Estes et al. 2011, Bailey et al. 2010). Our observations are consistent with those of Shackell et al. (2010), who demonstrated that due to fishing, predator evolution toward smaller sizes off the western Scotian Shelf led to a 300% increase in prey biomass (Shackell et al. 2010). While the consequences observed in our study are more nuanced (0-2% increase in prey biomass), they nevertheless underscore the broader implications of these evolutionary changes, particularly through the generation of trophic cascades (Shackell et al. 2010).

FIE has been suggested to potentially threaten fishery yields (Conover and Munch, 2002) or hinder stock recovery once fishing ceases (Olsen et al. 2004; Hutchings 2000; de Roos et al. 2006). Our results support this view, as we show that fishing yields are generally reduced by body size evolution. While prey evolution almost invariably leads to reduced yields, predator evolution toward larger sizes may increase yields. However, these outcomes are only observed when fishing targets smaller predators, which, through evolution, become less susceptible to capture. Evolution thus promotes yields by lowering the fishing pressure experienced and allowing predators to grow larger. Such strategies are rarely employed, as fisheries generally target large individuals (Jorgensen et al. 2007, Darimont et al. 2009), but align with the recommendations of Hixon et al. (2014), who advise against fishing large individuals (BOFFFFs, Hixon et al. 2014). Secondly, our findings reveal that evolution can also enhance the resilience of overexploited populations. By evolving toward smaller sizes, prey species can contribute to the prolonged persistence of their predators. In certain systems, the co-evolution of prey and predators enables the latter to survive disturbances that would otherwise drive them to extinction. This phenomenon of evolutionary rescue-both direct (predator evolution) and indirect (prey evolution) - has been documented (Vander Wal et al. 2012, Carlson et al. 2014, Yamamichi and Miner, 2015). Our study further suggests that it could support both the conservation of exploited species and the sustainability of the associated fisheries.

Using a simple system allows for an analytical understanding of the selection pressures generated by fishing. However, these simplifications come at the expense of certain approximations. From an ecological perspective, each species is represented by a single size, which fails to capture the diversity of ages and sizes within populations. Employing an age-structured model (such as that of Ernande et al. 2004) adapted to a community context could provide a more nuanced understanding of the direct and indirect effects of fishing on size evolution. Furthermore, we do not account for cannibalism, despite this phenomenon being recognized as significant and capable of altering the direction of natural selection (Edeline et al. 2007). Finally, our study does not fully capture the complexity of marine systems. Complementary approaches, such as those based on interaction networks, would be necessary to generalize our findings (Hocevar et al. 2021, Loeuille, 2019).

From an evolutionary perspective, we initially employed adaptive dynamics to investigate the effects of fishing on size evolution. This approach assumes slow evolution and a clear separation between ecological and evolutionary timescales. While it provides a way to properly define fitness based on the eco-evolutionary loop, and here a qualitative understanding of the balance between direct and indirect effects of fishing, its assumptions contradict observations of rapid size evolution (Olsen et al. 2004). Law and Plank (2018) argue that such an approach is overly ambitious because fishing-induced disturbances are too intense to maintain a separation between ecological and evolutionary timescales. Nevertheless, other authors have justified the use of these tools for this type of analysis (see research groups led by Ulf Dieckmann and Miko Heino), and the third part of our study does not rely on this assumption. Finally, these types of models are poorly suited to real-world fishing conditions. We believe that if body sizes evolve, fishers will adjust their practices accordingly, and regularly modifying target sizes could further amplify selection effects on body size.

Despite these limitations, our study highlights several important points. First, community structure is critical for understanding size evolution, aligning with calls for ecosystem-based fishery management (Pikitch et al. 2004). Second, it demonstrates that predators are inherently more susceptible to direct selection, further emphasizing the need for their conservation. Finally, it reveals that selective forces can counteract each other, resulting in an apparent absence of evolution. While some advocate minimizing FIE (Kuparinen and Merila, 2007, Law and Plank, 2018), promoting such balancing mechanisms may nonetheless erode the future evolvability of populations (Edeline and Loeuille, 2021), potentially threatening their survival. We suggest that limiting these evolutionary changes could be achieved by distributing fishing-induced mortality across various species within ecosystems (Zhou et al. 2010, Garcia et al. 2012, Law and Plank, 2018) and, more broadly, by adopting a precautionary approach to fisheries management (Pikitch et al., 2004), including regular monitoring of exploited population traits (Conover and Munch, 2002).

## Supplementary materials

### Supplement 1: Life-history and interaction parameters

The life-history parameters and the strength of predatory interactions depend on relative prey and predator sizes (expressed as a log(biomass) *z*_*i*_). Mortality rate is assumed to be decreasing with size (Brown et al. 2004):

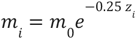

Attack rate is a function of predator-prey size ratio with a maximal value when predators are *d* times larger than their prey (Brose et al. 2006) :

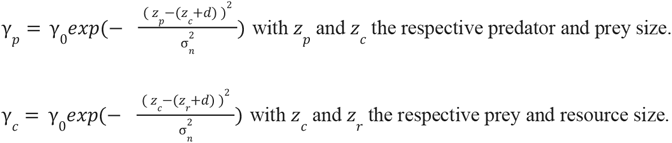

The conversion efficiency is the efficiency of converting one unit of prey *j* into one unit of predator *i* and decreases with size difference:

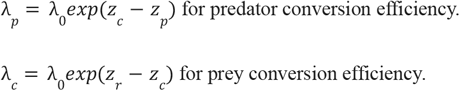

In this study, we consider resource size *z*_*r*_ = 1, and we assume preferred predator-prey size difference *d* = 5. 65 to be the same for consumers and predators. Competition is taken into account but is intraspecific and therefore does not depend on size differences. Other parameters values: σ_*n*_ = 1.

### Supplement 2: Ecological equilibria, feasibility and stability

The fishing intensity

*E*, the allocation of effort between the two species (*q*_*p*_, *q*_*c*_ with *q*_*p*_ + *q*_*c*_ = *q*_*tot*_) and the intraselectivities used (*s*_*p*_, *s*_*c*_), determine the persistence of populations under exploitation. Specifically, the Lotka-Volterra dynamic system (eq. 1, eq. 2) exhibits three distinct equilibria (*C*\*, *P*\*):

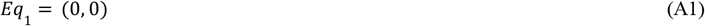

where both species go extinct due to fishing pressure,

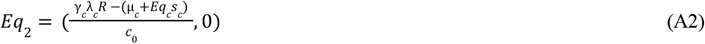

where predators are driven to extinction, while prey persist,

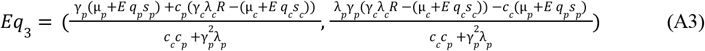

where fishing allows the coexistence of both species.

To analyze the feasibility and stability conditions of these equilibria, we define 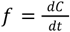 and 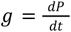 as the rate of change for prey and predator populations, respectively (eq. 1, eq. 2). The associated Jacobian matrix *J* is expressed as:

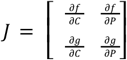

For each equilibrium, the Jacobian matrix *J* takes the following form:

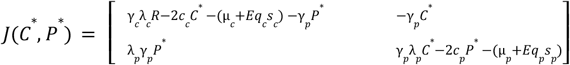

#### Analysis of equilibrium *Eq*_3_

We aim to determine the fishing conditions that allow the coexistence of prey and predators. For *Eq*_3_ to exist, both population densities must remain positive. Prey populations persist as long as *E* < *E*_*c*_, where:

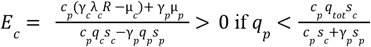

This critical threshold *E*_*c*_ represents the maximum fishing intensity beyond which prey populations cannot be sustained. Additionally, predator persistence requires that their population density remains above zero, which imposes further constraints on fishing strategies and effort distribution. When the effort distribution toward predators exceeds 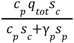, prey populations cannot go extinct. Predators, however, are maintained as long as *E* < *E*_1_, where:

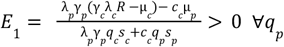

The difference *E*_*c*_ – *E*_*p*_ is positive if 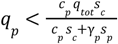. Specifically:

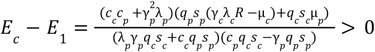

Thus, when *E* < *E*_1_, both prey and predators coexist in the system, making *Eq*_3_ feasible. For *Eq*_3_ to be stable, the trace (*tr*) and determinant (*det*) of the Jacobian matrix *J* at equilibrium *Eq*_3_ must be negative and positive respectively.

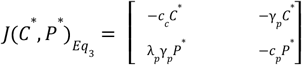

when *P*\* > 0 and *C\** > 0, 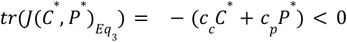

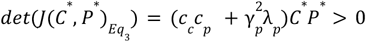

Therefore, when *P*^*^ > 0 and *C*^*^ > 0, in other words when *E* < *E*_1_, *Eq*_3_ is feasible and stable.

#### Analysis of equilibrium *Eq*_2_

The fishery maintains prey populations if the fishing effort does not exceed *E*_2_. Thus, when *E* < *E*_2_, *Eq*_2_ is feasible.

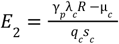

To assess the stability of this equilibrium, we examine the trace and determinant of the Jacobian matrix *J* at equilibrium *Eq*_2_ :

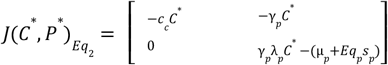

When *E* < *E*_2_, *C*\* > 0,

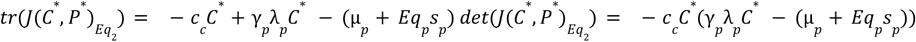

For *Eq*_2_ to be stable, the trace and determinant of *J* at *Eq*_2_ must be negative and positive, respectively. The feasibility conditions alone are not sufficient for the stability of *Eq*_2_. It is observed that:

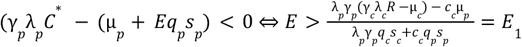

Therefore, when *E*_1_ < *E* < *E*_2_, *Eq*_2_ is stable.

#### Analysis of equilibrium *Eq*_1_

*Eq*_1_ = (0, 0) and is always feasible.

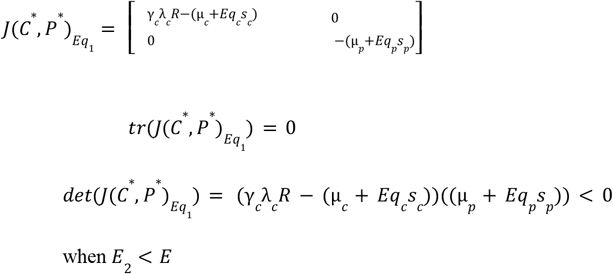

Thus, when the fishing effort exceeds *E*_2_, the system reaches *Eq*_1_ (a saddle point).

### Supplement 3: Ecological implications of fishing

Under coexistence conditions (see Supplementary S2), fishing affects prey (*C*^*^) and predators (*P*^*^) densities differently. From eq. A3 we can write:

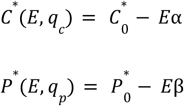

With 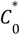 and 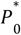 prey and predator densities without exploitation respectively. α and β capture the effect of the fishing pressure *E* on densities.

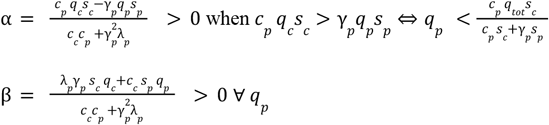

In other words,

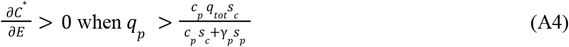

because prey benefit from predator removal

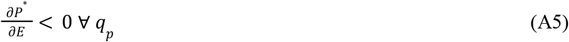

and predators are always negatively affected by fishing, because fishing both directly targets them and indirectly competes with them for prey.

### Supplement 4: Predator evolution and coevolution models (*b* and *c*)

#### Predator evolution only (model b)

Predator size (*z*_*p*_) evolution is assessed using the canonical equation of adaptive dynamics (eq. A6, Dieckmann and Law, 1996).

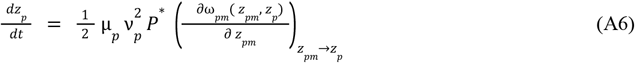

Where μ_*p*_ denotes the *per capita* mutation rate of predators and *ν*_*p*_ the mutation amplitude (see Table 1 in main text). ω_*pm*_ is called the invasion fitness (Metz et al. 1992), defined by the rate of variation of the density of rare mutants of body size *z*_*pm*_ in a resident population of trait *z*_*p*_ (eq. A7):

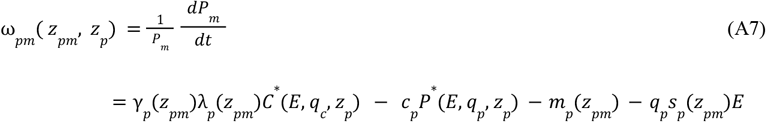

Prey body size is assumed not to evolve in this case (*b*). According to eq. A6, the evolution of predator size depends on the ability of nearby mutants to invade the system defined by residents. Evolution proceeds if the population persists (*P* * ≠0) and as long as proximate sizes confer a competitive advantage (∂ω_*pm*_ ≠0). When the selection gradient (eq. A8) is positive (negative), it means that a mutant of larger size has better (worse) fitness compared to the residents, and this is reflected in the canonical equation (eq. A6) by the increase (decrease) in size through invasion. Harvesting the community through interspecific (*q*_*p*_, *q*_*c*_) and intraspecific (*s*_*p*_) selectivities may affect predator evolution both indirectly, via changes in community structure (changes in *C*^*^ in first term of eq. A8), and directly, through selection on predator body size (third term of eq. A8):

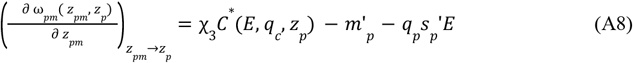

where

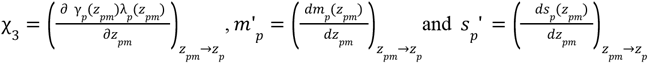

Depending on the sign of the first and third terms of eq. A8, DS and IS may either act in the same or in opposite directions, thereby influencing body size evolution. Once the selection gradient vanishes, the evolution of predator size stops and reaches the evolutionary singularity 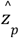.

#### Coevolution (model c)

Predator and prey coevolution is assessed using the canonical equation of adaptive dynamics (eq. A9&A10).

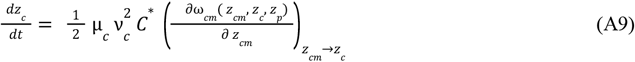

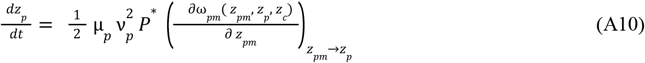

Where the fitnesses ω_*cm*_ and ω_*pm*_ are:

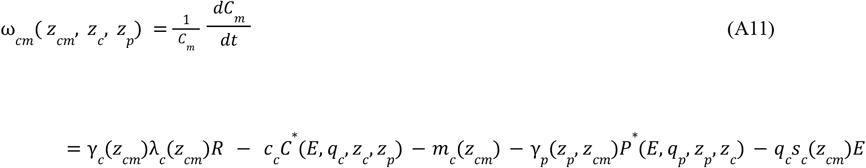

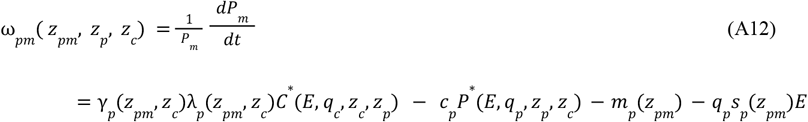

Harvesting the community through interspecific (*q*_*p*_, *q*_*c*_) and intraspecific (*s*_*p*_, *s*_*c*_) selectivities may affect prey and predator evolution both indirectly, via changes in community structure (changes in *P*^*^ and in *C*\* in first and third term of eq. A13 and A14 respectively), and directly, through selection on the body size of both species (last term of eq. A13 and A14 for prey and predator respectively):

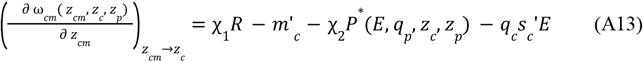

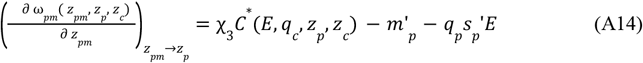

where

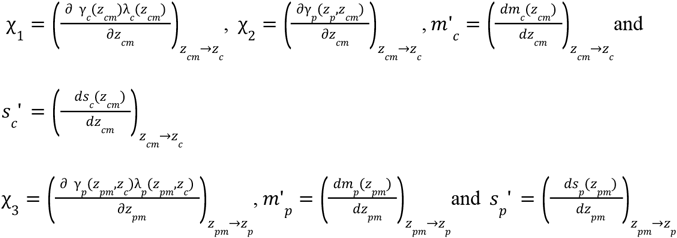

In this case, altering community structure affects the trophic constraints on prey and predator body sizes through the terms χ_2_ *P* * (*E, q*_*p*_, *z*_*c*_, *z*_*p*_) for prey and χ_3_ *C* (*E, q*_*c*_, *z*_*p*_, *z*_*c*_) for predators. In each of the equations (eqs. A13 & A14), DS and IS may act either in the same or in opposite directions, thereby influencing body size evolution. Once both selection gradients vanish, coevolution ceases and sizes reach their evolutionary singularities, 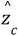 and 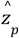.

## Supplement 5: Size evolution in response to IS

Since the reasoning is analogous for the other models, we only detail the demonstration for model *a*. The demonstration for model *b* can be obtained by setting θ_*p*_ = 0 in the IS + DS demonstration provided in Supplementary S6 (eq. A26).

At eco-evolutionary equilibrium, in the absence of DS (θ_*c*_ = θ_*p*_ = 0), we obtain:

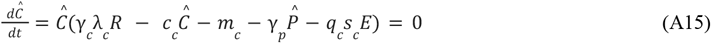

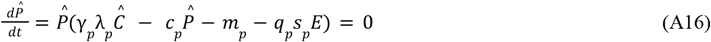

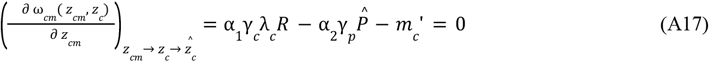

with 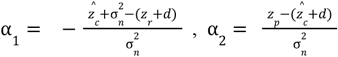 and 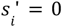 without DS.

eq. A17 defines the conditions under which the fitness gradient vanishes, thereby determining the evolutionary singular strategy of prey body size 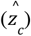. Once this strategy is reached, the corresponding ecological equilibrium densities of prey and predators are given by *Ĉ* and 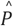 respectively. To understand how fishing affects the evolution of prey body size through eco-evolutionary feedbacks, we examine how an increase in fishing effort modifies the evolutionary singularity condition (eq. A17) together with the ecological equilibrium conditions for prey and predator densities (eqs. A15 and A16). Implicit differentiation of eq. A17 with respect to fishing effort, accounting for its effect on prey size, yields:

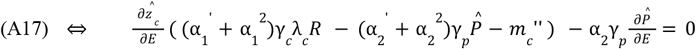

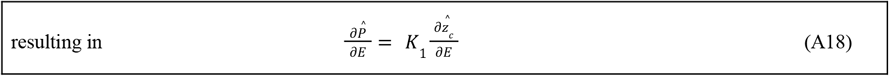

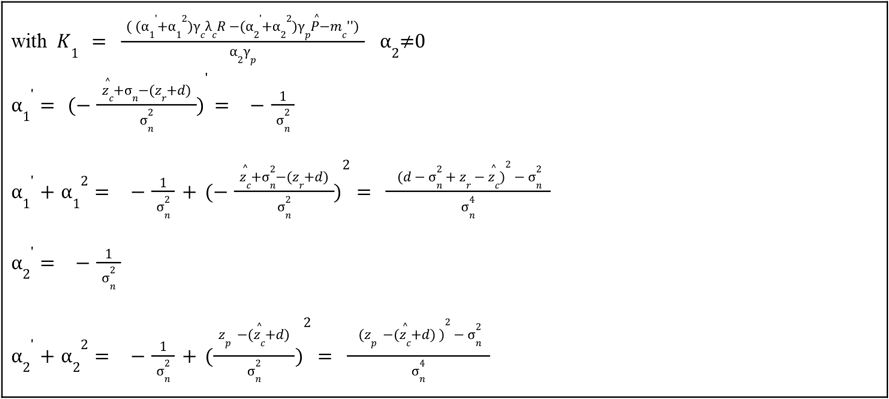

This decomposition (eq. A18) shows that, in response to increased fishing pressure, prey body size would evolve in the same direction as changes in predator densities if *K*_1_ is positive and in the opposite direction if *K*_1_ is negative.

We now consider the effect of increasing fishing pressure on the ecological equilibrium condition governing prey (eq. A15) and predator dynamics (eq. A16) using implicit differentiation:

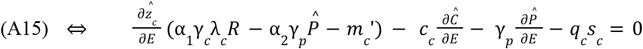

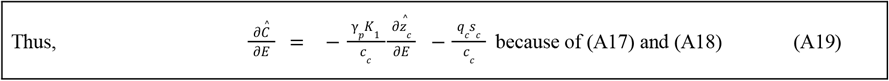

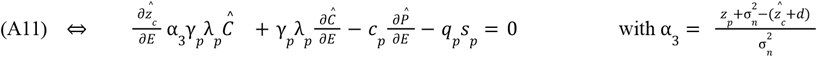

According to (A18) and (A19),

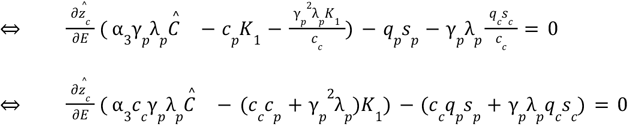

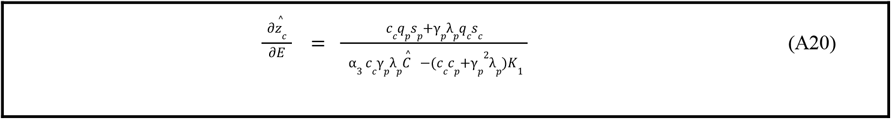

Because the numerator of eq. A20 is strictly positive, fishing always induces prey body size evolution in absence of direct selection. Note that this result does not hold true for predator size evolution (*model b*, setting θ_*p*_ = *s*_*p*_′ = 0 in eq. A26 from Supplementary S6), because 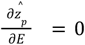 when *q*_*p*_ *s*_*p*_ *γ*_*p*_ = *q*_*c*_ *s*_*c*_ *c*_*p*_. This difference (between A20 and A26 when setting *s*_*p*_′ = 0) arises because in absence of direct selection, the only driver of a species’s evolution are the changes in the other specie’s densities. Because prey densities can either increase or decrease with an increased fishing pressure (Supplementary S3), predator body size evolves accordingly. On the opposite, predators only suffer from an increased fishing pressure (Supplementary S3) thereby always inducing prey downsizing.

We now investigate whether fishing induces an evolutionary increase or decrease of prey body size. Before fishing starts, α_3_ is close to 0 because 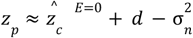 (at eco-evolutionary equilibrium without fishing, predators are close to the optimal size for their resource acquisition). In addition, using the non-invasibility criterion of ESS (Metz et al., 1992) we obtain from eq. A17:

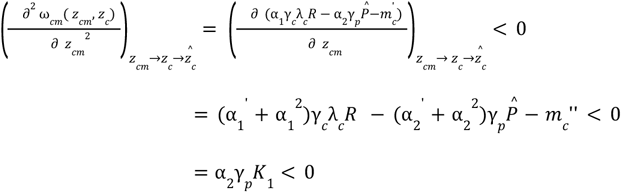

Because 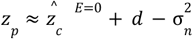, at low exploitation levels 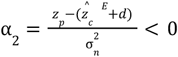 and thus *K*_1_> 0.

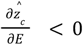

Thus,

This reasoning is valid only for low levels of exploitation, but it provides insight into the direction of prey size evolution in response to fishing.

## Supplement 6: Size evolution in response to IS + DS

We focus here on predator evolution under both types of selection, while the results for prey evolution are provided at the end of this section (eq. A28). For predator evolution, we obtain, assuming convergence to eco-evolutionary equilibrium:

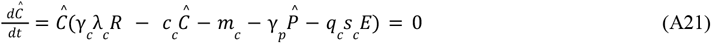

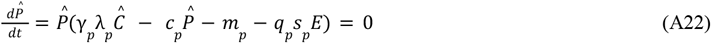

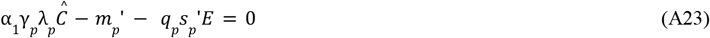

with 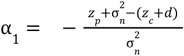 and 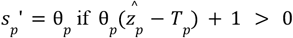 (eq. 3 from main text).

eq. A23 defines the conditions under which the fitness gradient vanishes, thereby determining the evolutionary singular strategy of predator body size 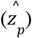. Once this strategy is reached, the corresponding ecological equilibrium densities of prey and predators are given by *Ĉ* and 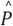 respectively. To understand how fishing affects the evolution of predator body size through eco-evolutionary feedbacks, we examine how an increase in fishing effort modifies the evolutionary singularity condition (eq. A23) together with the ecological equilibrium conditions for prey and predator densities (eqs. A21 and A22). Implicit differentiation of eq. A23 with respect to fishing effort, accounting for its effect on predator size, yields:

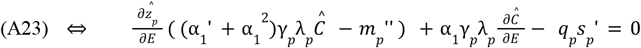

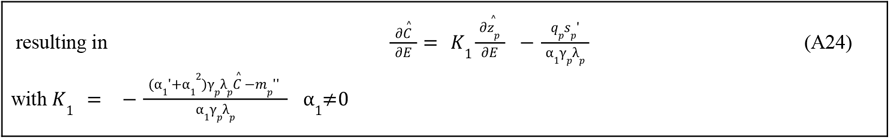

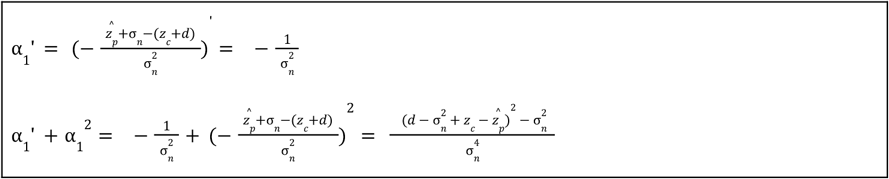

We now consider the effect of increasing fishing pressure on the ecological equilibrium condition governing prey (eq. A21) and predator dynamics (eq. A22) using implicit differentiation.

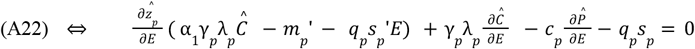

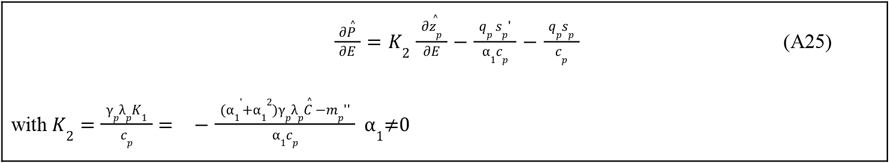

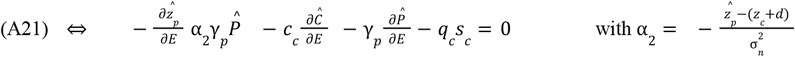

Using eq. A24 & A25,

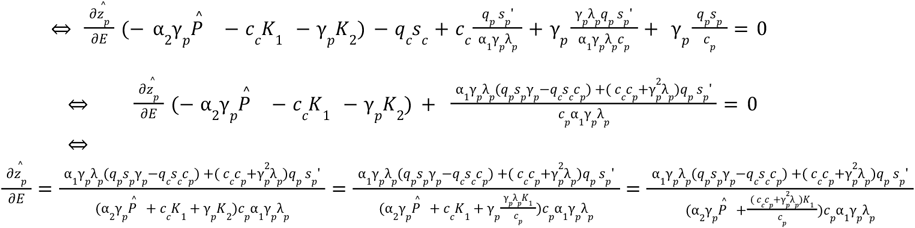

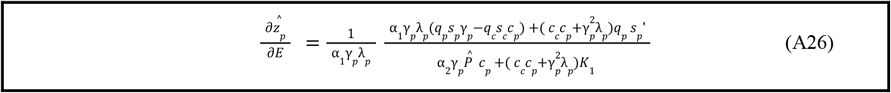

Note that this equation (A26) still hold true when we only consider IS and remove DS *s*_*p*_′ = 0). Note also that since prey do not evolve in this model (*b*), *s*_*c*_ = 1 because *z*_*c*_ = *T*_*c*_ (eq. 3 from main text).

To find the sign of 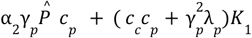, we can show that

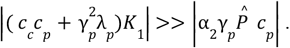

Using the non-invasibility criterion and a reasoning analogous to that in Supplementary S5, we can show that α_1_ < 0 and therefore *K*_1_ < 0.

Thus,

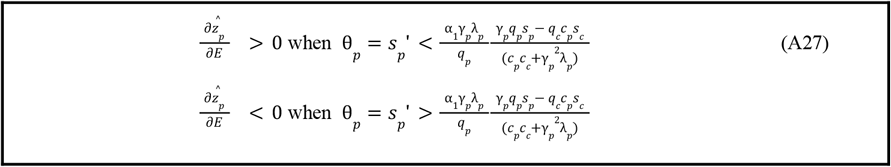

For prey evolution, we have:

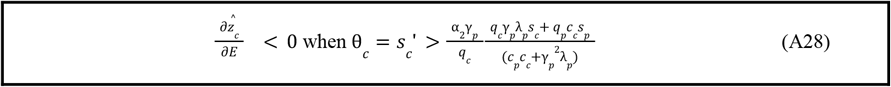

### Supplement 7: Sensitivity of prey evolution to initial size distribution

**Figure S7:**
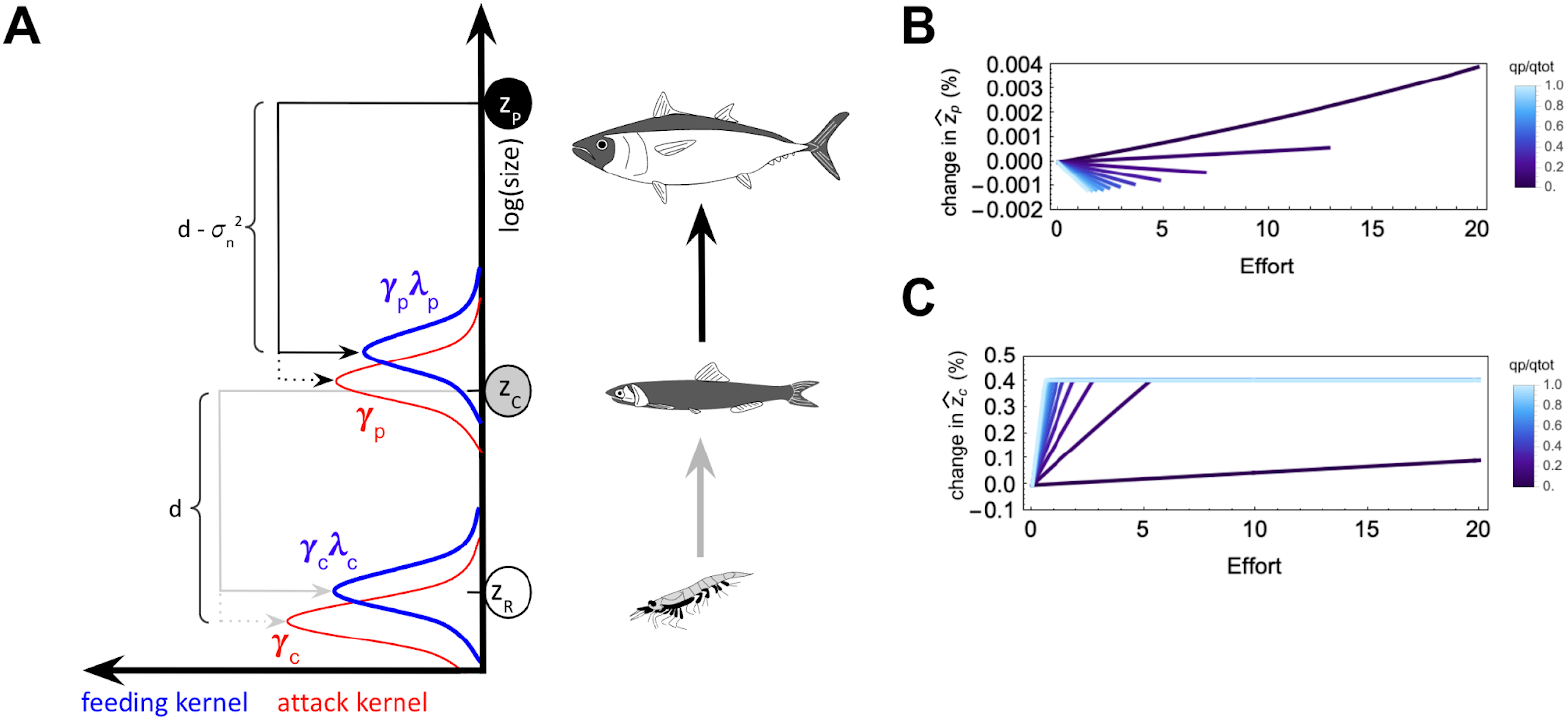
Effects of fishing on body size evolution in the absence of DS and when species are not at eco-evolutionary equilibrium prior to fishing. A: Community and associated trophic interactions illustrating a scenario away from eco-evolutionary equilibrium. B: evolution of predator body size (model *b*) without DS and under different distributions of fishing effort. C: evolution of prey body size (model *a*) without DS and under different distributions of fishing effort.

When prey and predator sizes are not initially at eco-evolutionary equilibrium (Figure S7A), fishing induces the same body size evolution in predators (Figure S7B) but an opposite pattern to that described in the main text for prey evolution (Figure S7C). Indeed, when only predators are allowed to evolve, their body size changes to optimize resource acquisition, resulting in the same situation as in Figure 2. Conversely, when only prey evolve and predators exhibit peak predation on sizes larger than those of the prey, we initially observe:

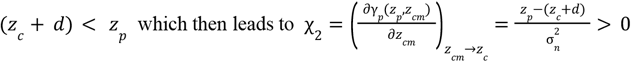

Thus, in this situation predators keep prey small, and their decline due to fishing promotes the evolution of prey toward larger body sizes 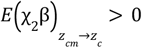 in eq. A29 and ∂ ω_*cm*_ ^*E*=0^ is a decreasing function due non-invasibility criterion, Metz et al., 1992).

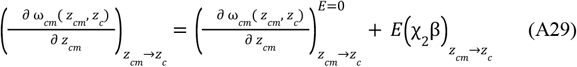

### Supplement 8: Size co-evolution in response to fishing (IS + DS)

We now examine the co-evolution of predator and prey body sizes in response to fishing, accounting for both types of selection.

#### Prey

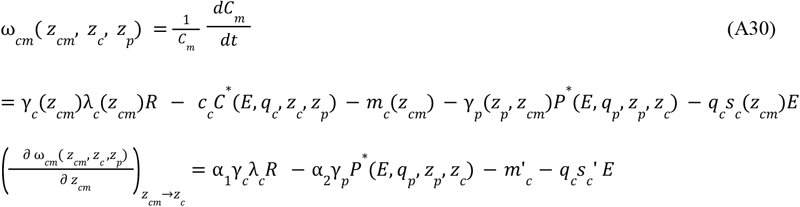

with 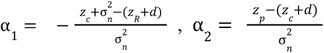

#### Predators

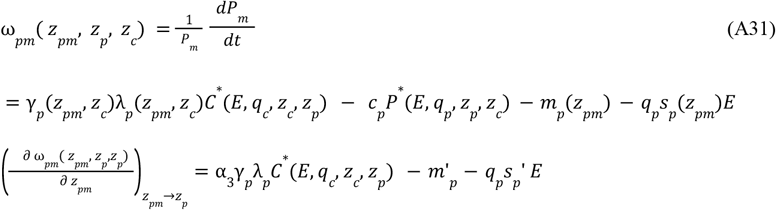

with 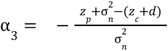

We then have the following system at eco-evolutionary equilibrium:

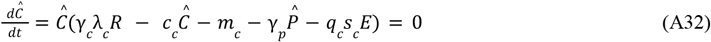

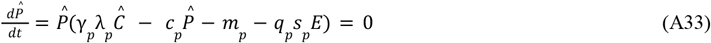

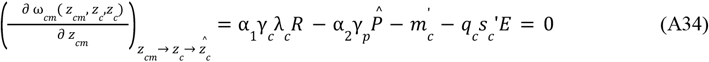

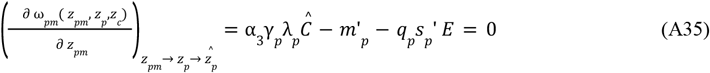

From eq. A35 in 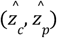 we have:

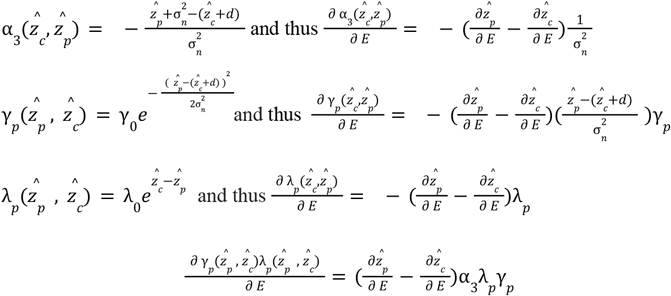

Remind eq. A35:

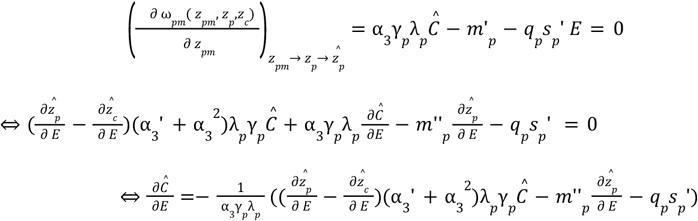

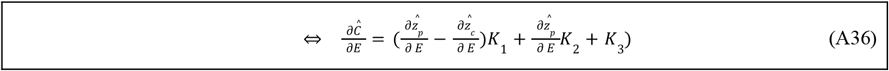

with 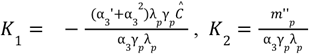 and 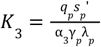

From eq. A34 in 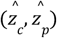 we have:

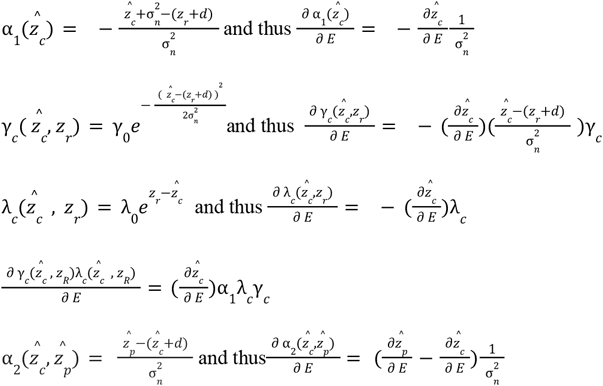

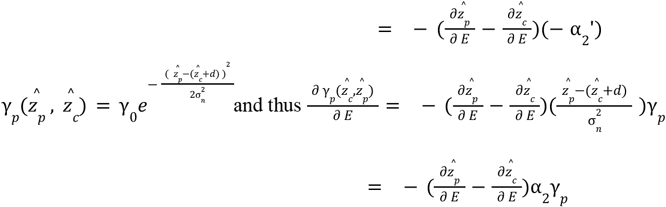

Remind eq. A34 :

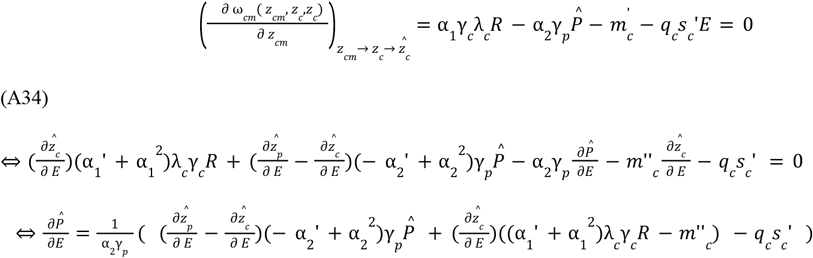

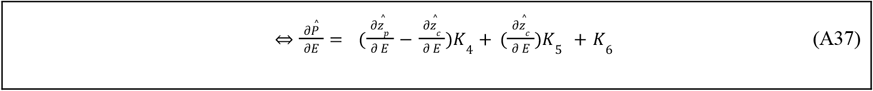

With 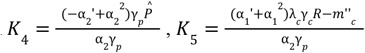 and 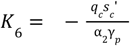

Remind eq. A32:

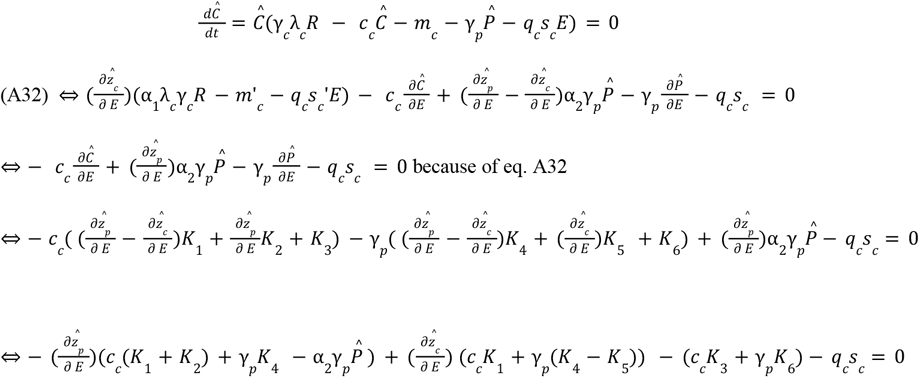

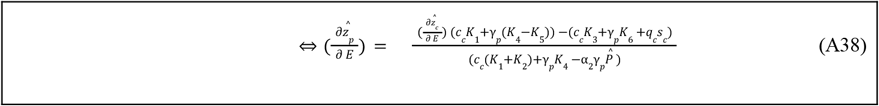

Remind eq. A33:

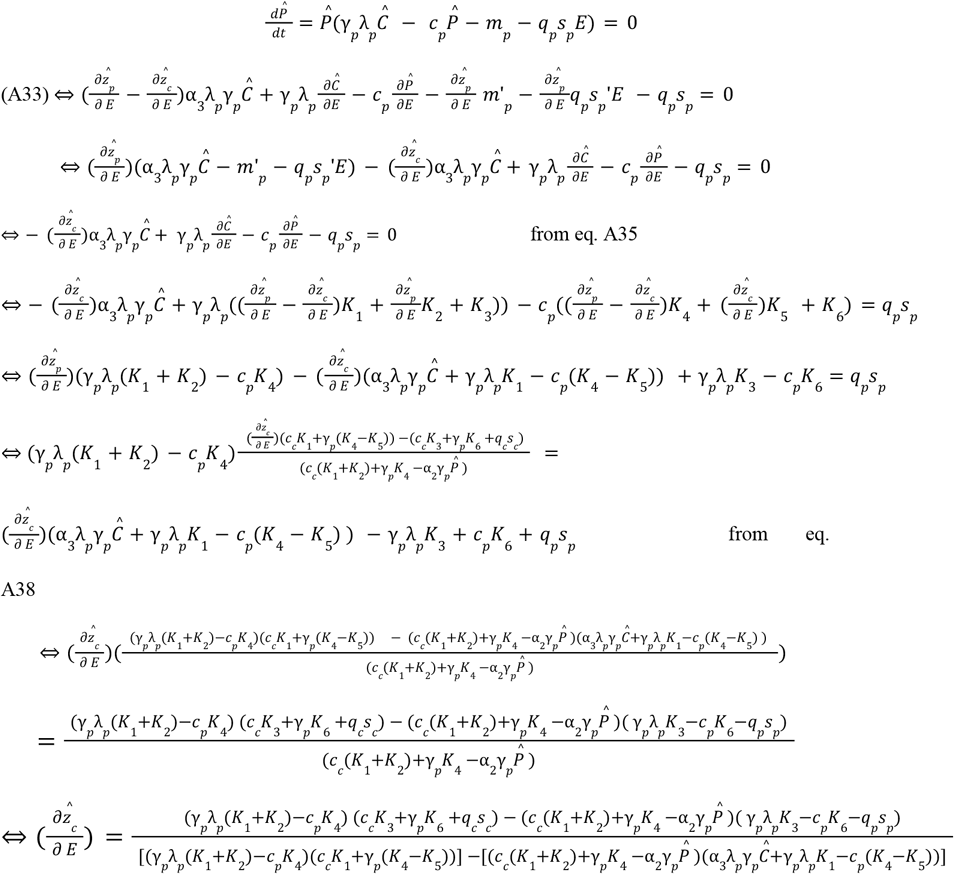

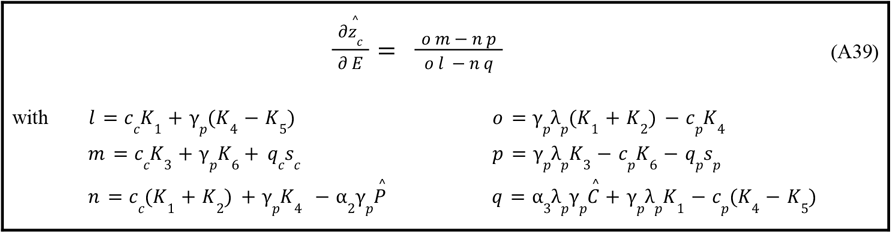

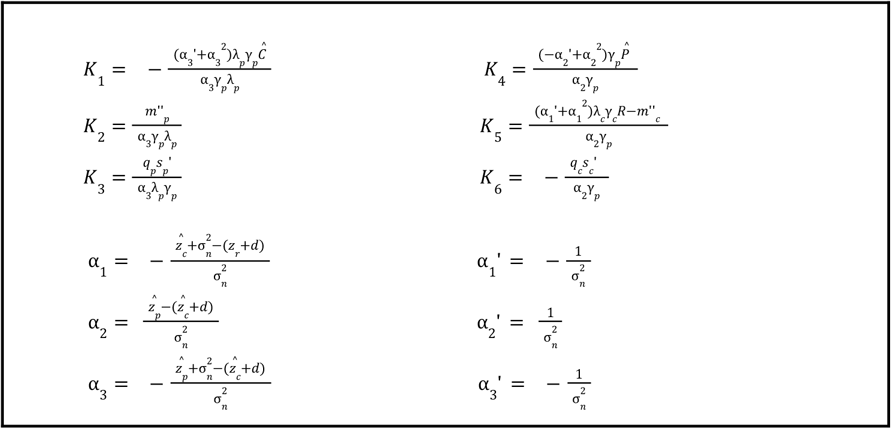

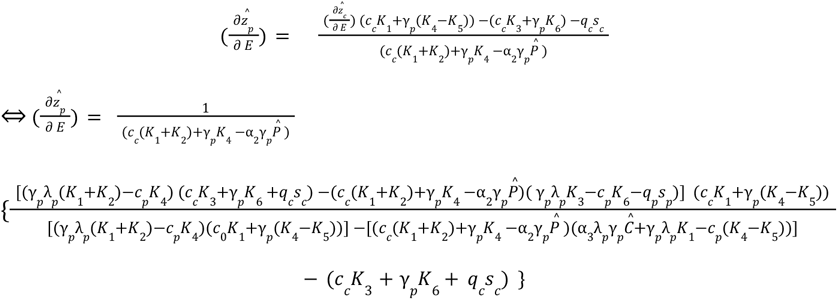

and because 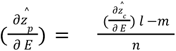

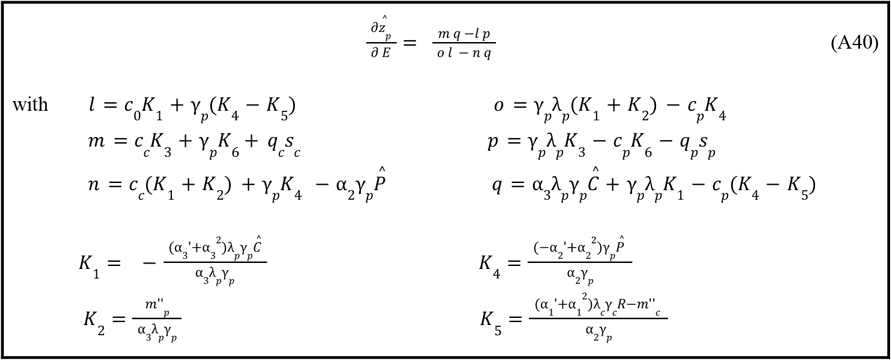

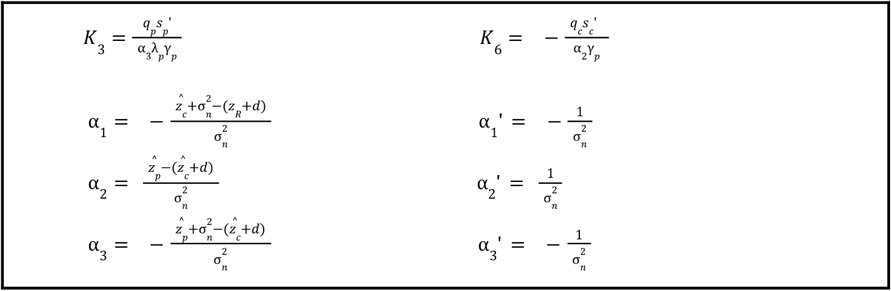

### Supplement 9: No evolution in response to fishing (IS + DS)

1. *We first seek management conditions on (s*_*c*_′, *s*_*p*_ ′, *q*_*p*_ *) such that there is no evolution of prey and predator body sizes when both species co-evolve*. Intuitively from eq. A37 we have,

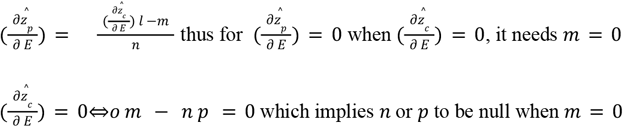

We note that *n* does not contain terms directly related to selectivities (i.e., to the slopes of the selectivity functions *s*_*p*_ and *s*_*c*_). Indeed, when 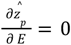 and 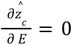 the terms *s*_*p*_ and *s*_*p*_ both equal 1 since 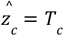 and 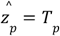. The question to address is therefore equivalent to:

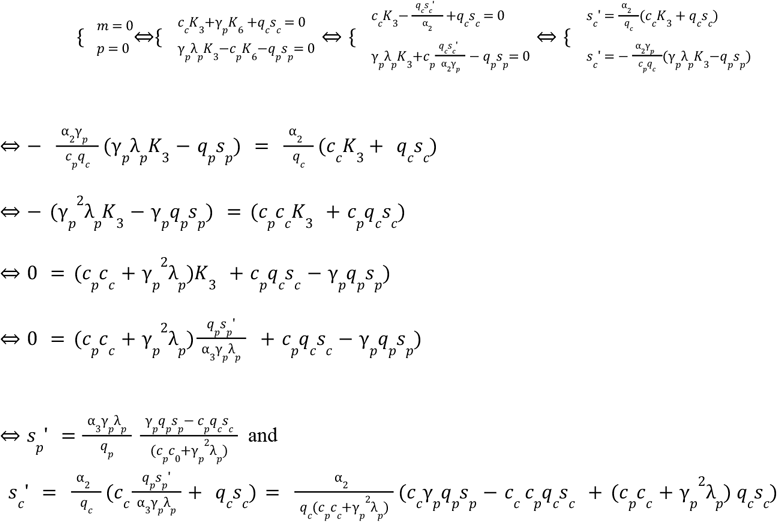

which leads to

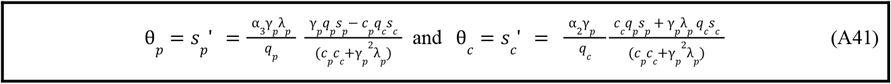

We note that these constraints are the same as when only a single species is allowed to evolve (considering that the size of the other species does not evolve, see Supplementary S6).
2. *We now seek management conditions on (s*_*c*_′, *s*_*p*_′, *q*_*p*_ *) such that prey body size cannot evolve when both species co-evolve*.

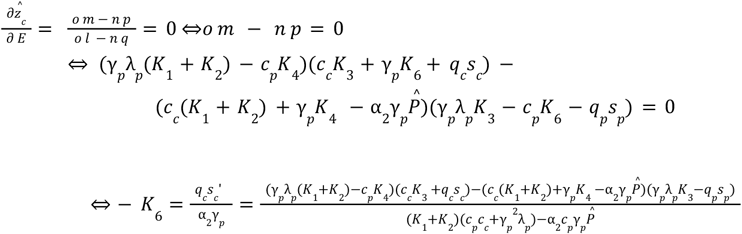

Thus, for all possible evolution of predator body size:

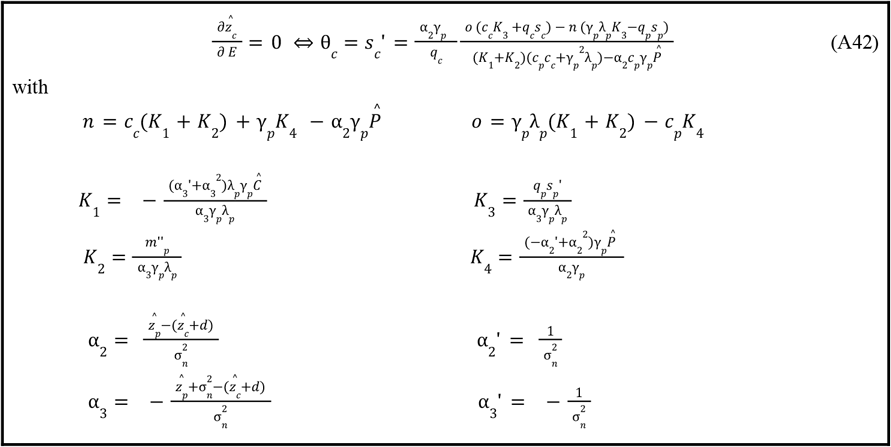

This result cannot be used as such because the α_*i*_ values are obtained at eco-evolutionary equilibrium. If 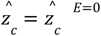 by definition, the 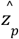 values are unknown. In other words, one cannot predict *s*_*c*_′ such that 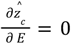 without an explicit formulation for 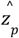.
3. *We now seek management conditions on (s*_*c*_′, *s*_*p*_ ′, *q*_*p*_ *) such that predator body size cannot evolve when both species co-evolve*.

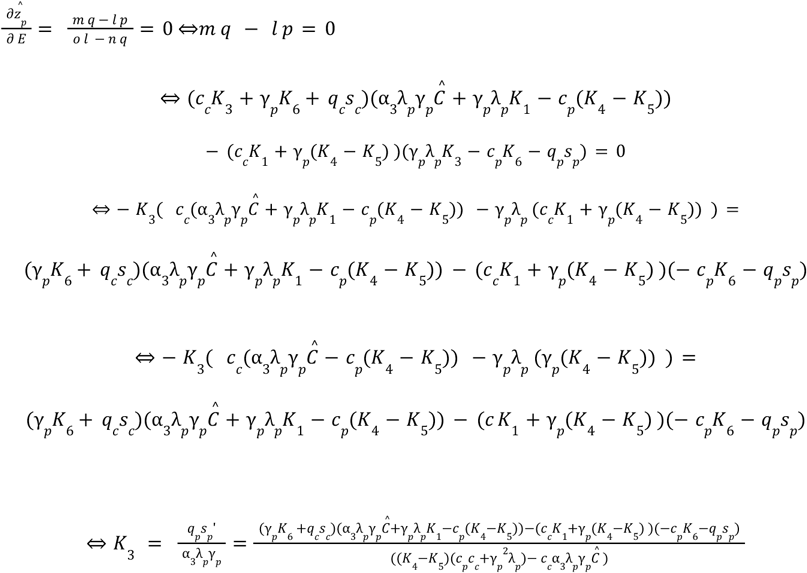

Thus, for all possible evolution of prey body size,

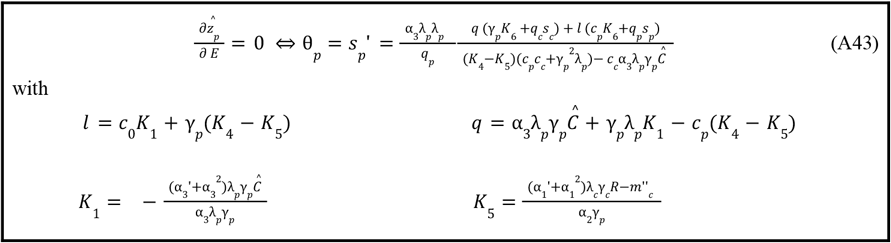

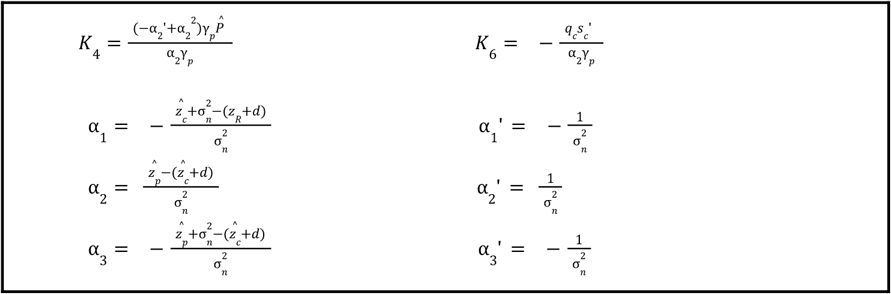

Again, this result cannot be used as such because the α_*i*_ values are obtained at eco-evolutionary equilibrium. If 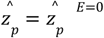 by definition, the 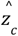 values are unknown. In other words, one cannot predict *s*_*p*_′ such that 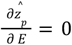 without an explicit formulation for 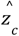.

### Supplement 10: Sensitivity of evolution to initial size distribution (IS + DS)

**Figure S10:**
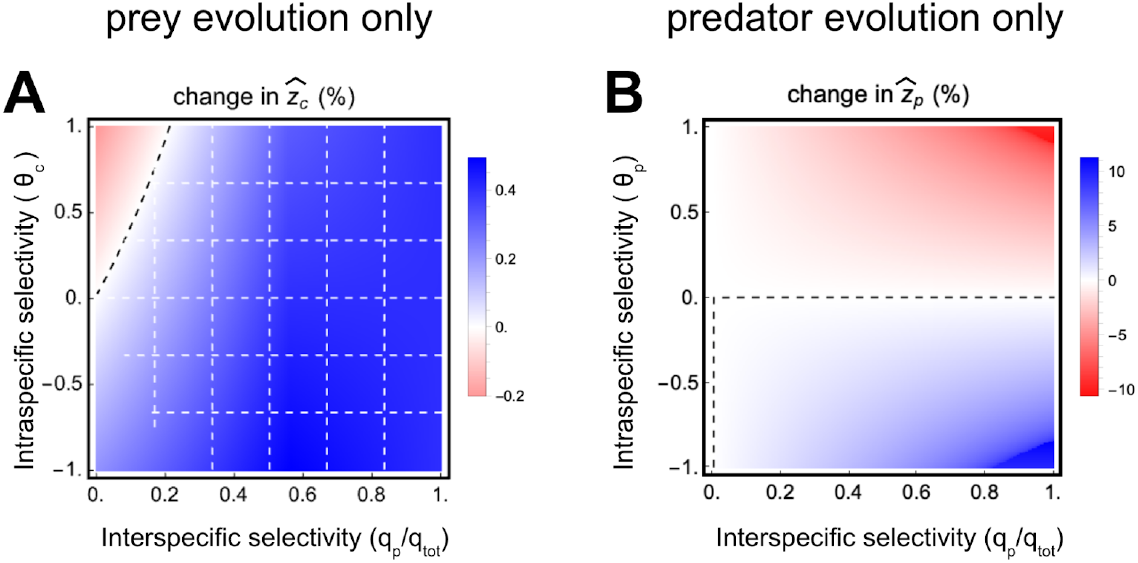
Evolution of sizes as a function of intraspecific and interspecific selectivity when only one species evolves. In this system, predator size is not initially at eco-evolutionary equilibrium (higher size, see Figure S1), which leads to an indirect selection which favors larger prey sizes. Depending on the nature of the fishing pressure, the size 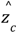 of preys (A) or 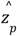 of predators (B) may either increase (in blue) or decrease (in red) due to evolution.The contribution of IS to size evolution can exceed that of DS (dashed white grid). DS and IS can offset each other, leading to the visible lack of size evolution (dashed black line). The fishing effort equals *E* = 1.

### Supplement 11: FIE (DS+IS) on prey body size and consequences on biomasses and community structure

**Figure S11:**
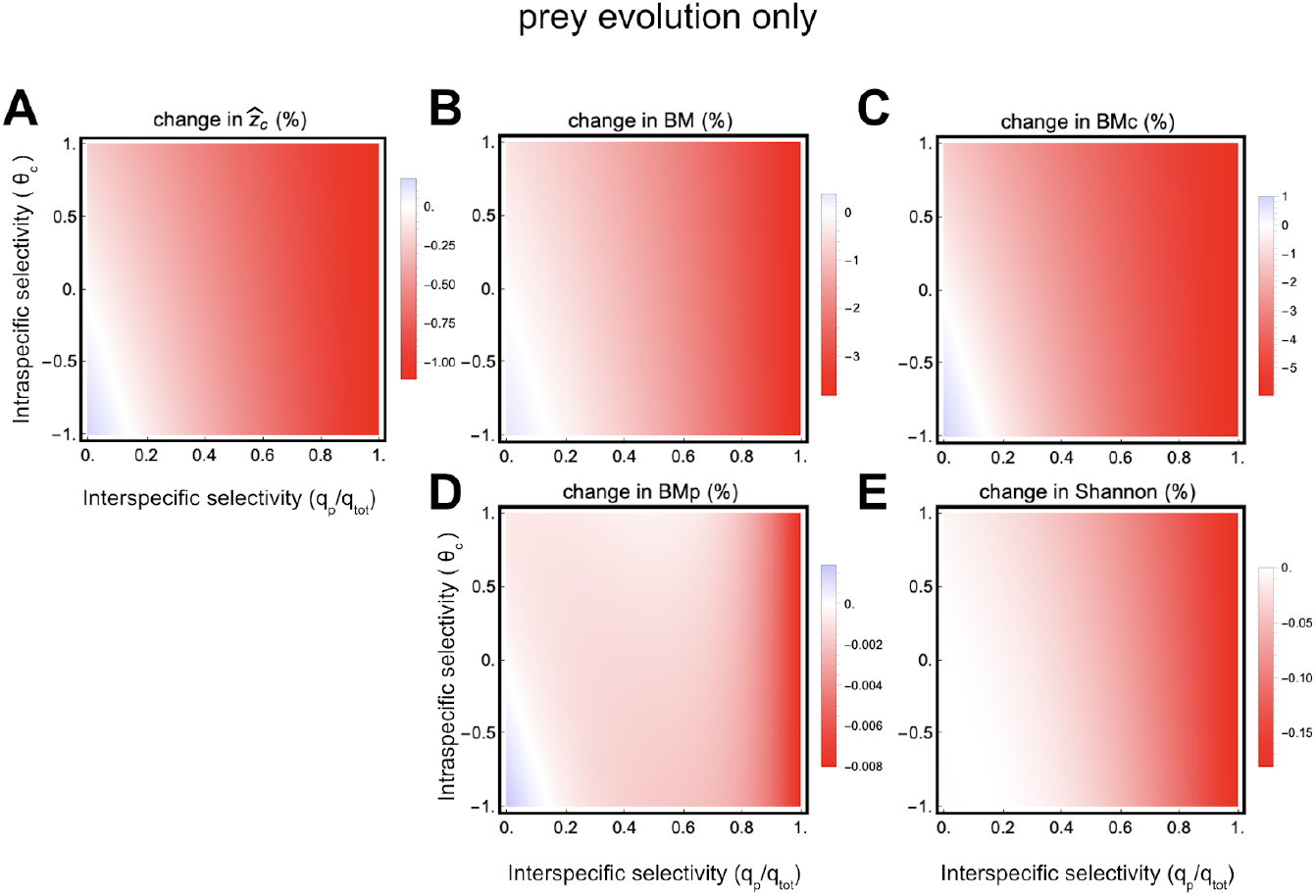
Consequences of prey size evolution on fish biomasses and density distribution within the community. Depending on the nature of the fishing pressure, the prey size 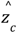, their biomass *BM*_*c*_, predator biomass *BM*_*p*_, total biomass *BM*, and Shannon index may increase (in blue) or decrease (in red) relative to a situation without evolution. θ_*c*_ is the slope of the intraspecific selectivity of the fishing effort targeting prey. *q*_*p*_/*q*_*tot*_ represents the share of the total effort *E* associated with fishing predators. The fishing effort equals *E* = 1.

### Supplement 12: FIE (DS+IS) on predator body size and consequences on biomasses and community structure

**Figure S12:**
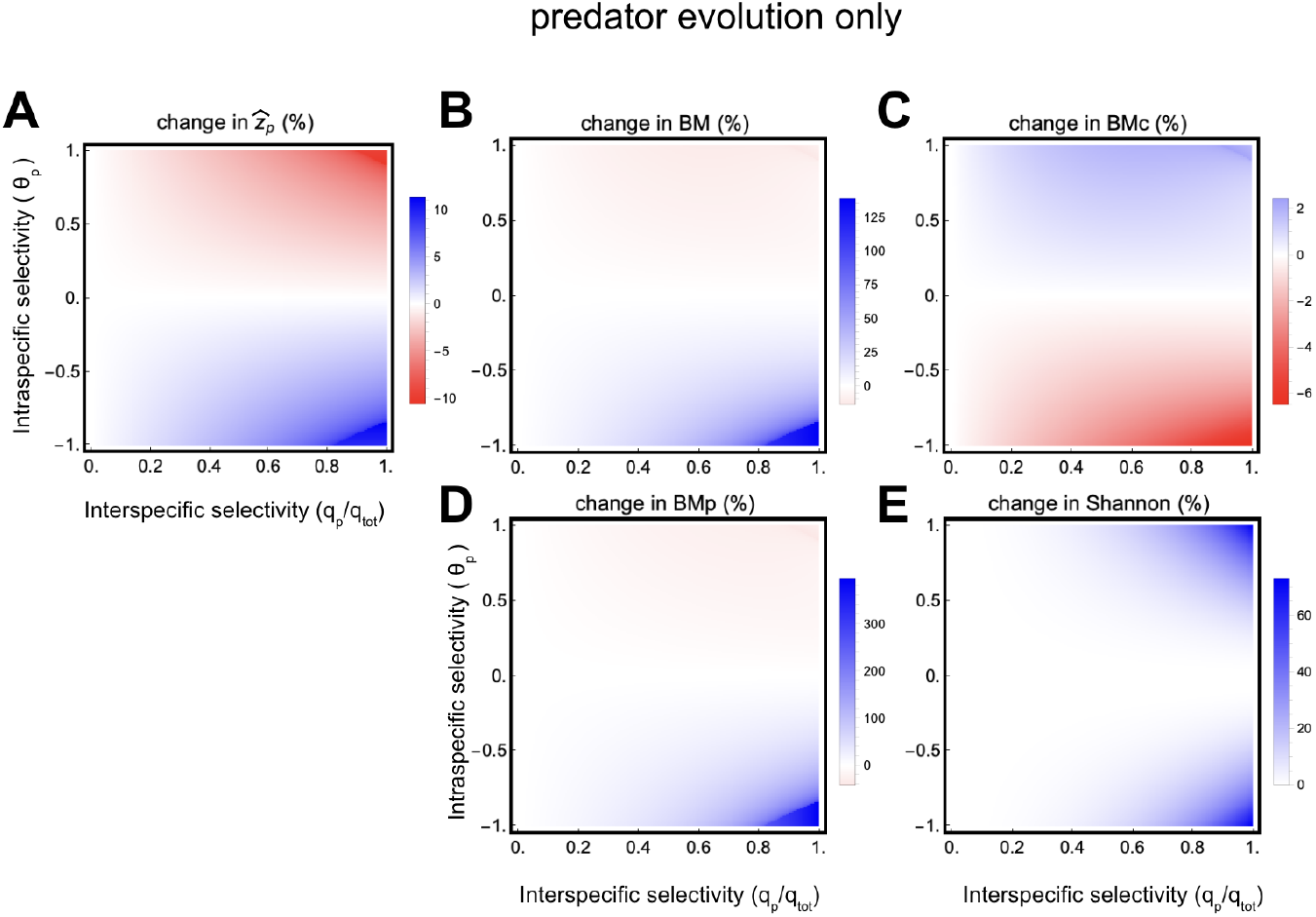
Consequences of predator size evolution on fish biomasses and density distribution within the community. Depending on the nature of the fishing pressure, the predator size 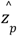, their biomass *BM*_*p*_, prey biomass *BM*_*c*_, total biomass *BM*, and Shannon index may increase (in blue) or decrease (in red) relative to a situation without evolution. θ_*p*_ is the slope of the intraspecific selectivity of the fishing effort targeting predators. *q*_*p*_/*q*_*tot*_ represents the share of the total effort *E* associated with fishing predators. The fishing effort equals *E* = 1.

### Supplement 13: FIE (DS+IS) on coevolving prey and predator body sizes

**Figure S13:**
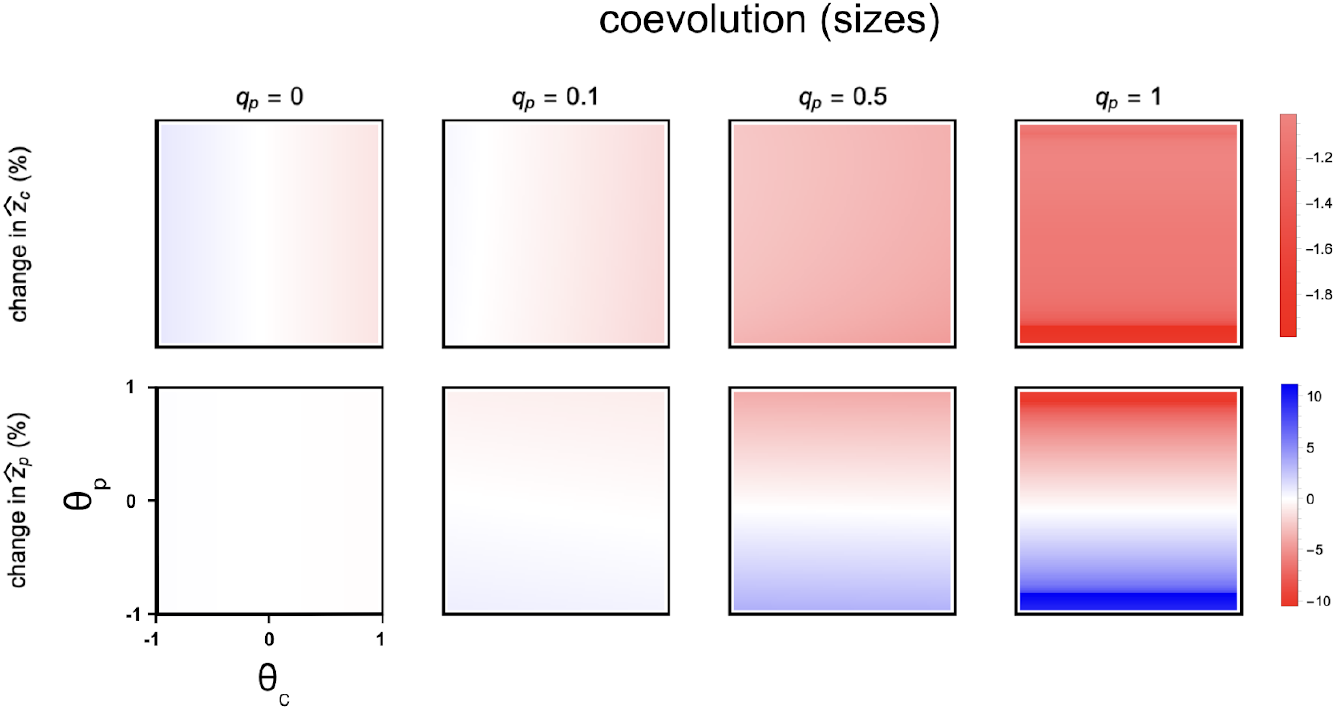
Co-evolution of prey and predator sizes depending on the nature of the fishing pressure. The size 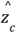 of preys and 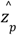 of predators co-evolve and may increase (in blue) or decrease (in red). θ_*c*_ and θ_*p*_ are the slopes of the intraspecific selectivity of the fishing efforts targeting prey and predators, respectively. *q*_*p*_/*q*_*tot*_ represents the share of the total effort *E* associated with fishing predators. The fishing effort equals *E* = 1.

### Supplement 14: FIE (DS+IS) on coevolving prey and predator densities

**Figure S14:**
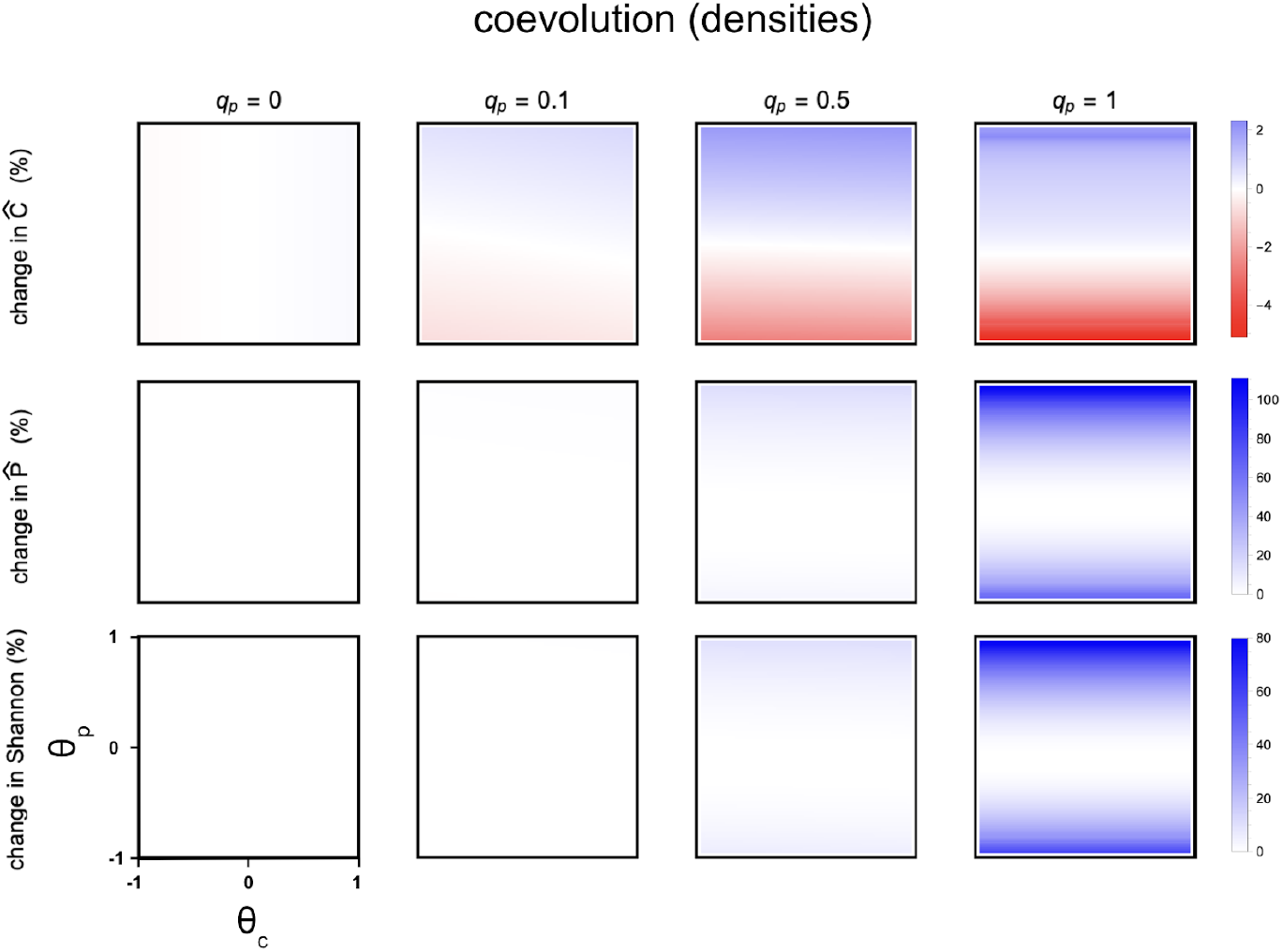
Consequences of the co-evolution of prey and predator sizes on their densities and distribution within the community. Depending on the nature of the fishing pressure, the densities *Ĉ* of the prey, 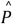 of the predators, and their distribution within the community may increase (in blue) or decrease (in red) compared to a situation without evolution. θ_*c*_ and θ_*p*_ are the slopes of the intraspecific selectivities of the fishing efforts targeting preys and predators, respectively. *q*_*p*_/*q*_*tot*_ represents the share of the total effort *E* associated with fishing predators. The fishing effort equals *E* = 1.

### Supplement 15: FIE (DS+IS) on coevolving prey and predator, biomass and yield

**Figure S15:**
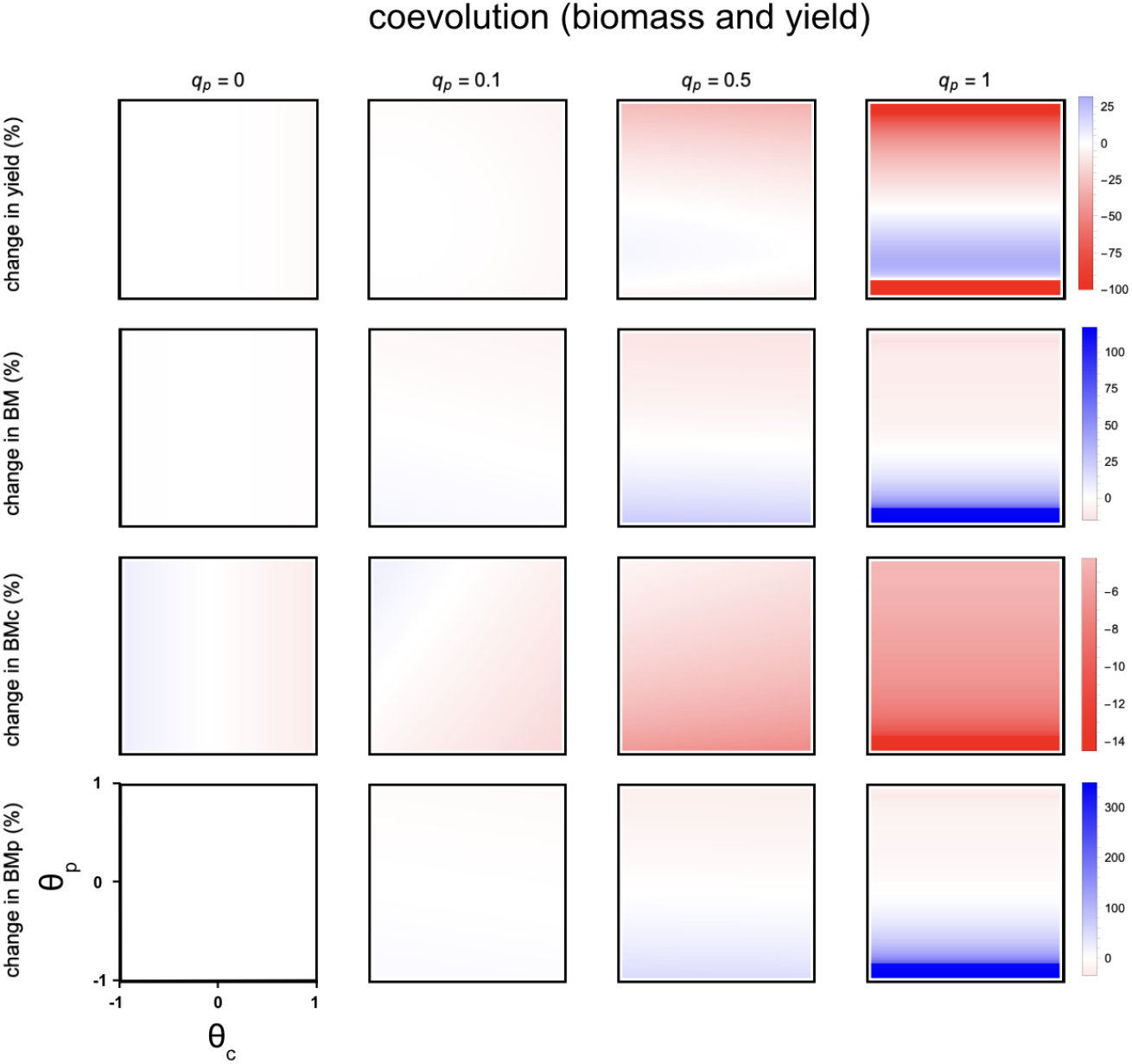
Consequences of the co-evolution of prey and predator sizes on yields and biomasses within the community. Depending on the nature of the fishing pressure, yields, the biomass of prey *BM*_*c*_, the biomass of predators *BM*_*p*_, and the total biomass *BM* within the community may increase (in blue) or decrease (in red) compared to a situation without evolution. θ_*c*_ and θ_*p*_are the slopes of the intraspecific selectivities of the fishing efforts targeting preys and predators, respectively. *q*_*p*_/*q*_*tot*_ represents the share of the total effort *E* associated with fishing predators. The fishing effort equals *E* = 1.

